# Hierarchical Clustering and Omics Analysis Revealed Massive Molecular Shifts in a Tree under Combined Hypoxia and Salinity

**DOI:** 10.1101/2023.03.24.534131

**Authors:** El-Hadji Malick Cisse, Bai-Hui Jiang, Li-Yan Yin, Ling-Feng Miao, Jing-Jing Zhou, Francine Ngaffo Mekontso, Da-Dong Li, Li-Shan Xiang, Fan Yang

## Abstract

Field and Greenhouse studies that attempted to describe the molecular responses of trees under waterlogging (WL) combined with salinity (ST) are quasi-inexistent. To dissect plant-specific molecular responses and patterns under SWL, we integrated transcriptional and metabolic analyses involving common differentially expressed genes (DEGs) and metabolites (DMs) patterns in *Dalbergia odorifera* leaflets. SWL-treated seedlings exhibited an impressively high number of DEGs and DMs compared to ST or WL. Although the main common DEGs and DMs showed a neutral pattern, gene ontology enrichment following the classification in different functional categories of SWL-transcripts displayed a predominant synergistic pattern mode. Hierarchical and silhouette analysis regrouped different morpho-physiological clusters following the treatment, SWL was mainly grouped with both single stress. SWL induced a massive shutdown of the photosynthesis apparatus through *LHCBs*- and *PSA*-related genes. Starch and plastoglobuli synthesis appeared to be drastically affected by SWL, while ABA content confirmed by ABA synthesis related-genes (*ABF*, *ABA1* and *NCED*) variations showed a less-needed role. *NXN* (Nucleoredoxin) genes are the main factors that sustain the antioxidant system under SWL. Here we provide the first molecular responses, characterization and patterns of a tree under SWL that would significantly shed light on our understanding of the molecular mechanisms underlying combined stress.

## Introduction

“The road is long, and the time is short, so we better be in a hurry”, was an acknowledgment by (Zandalinas and Mittler, 2022) in a recent report apropos of the lack of data involving multi-factorial stress combination effects on plant growth and development. It has been unanimously well established that human activity altered forest ecosystems and plant biodiversity by intensifying diverse environmental factors that tend to combine and cause more damage to tree species in their natural habitats. Many landscapes are exposed to synchronized waterlogging (WL) and salinity (ST) throughout the world. Indeed, the occurrence of hypoxia and ST simultaneously is growing in landscapes such as semiarid regions (Carter et al., 2006; Duhan et al., 2019). The earlier study focused on crops highlighted that the molecular response of plants under a combination of stresses is unequaled to one of the stress applied individually (Mittler, 2006). However, little is known about the molecular mechanisms underlying the responses of plant species to a combination of different stresses (Sewelam et al., 2020; Zandalinas et al., 2020), particularly in forest trees. Indeed, we do not know what tree species’ transcriptional and metabolomic variations hold to defend themselves against a combination of hypoxia and saline conditions. The limited pieces of information about the response of trees to a combined of WL and ST (SWL) concern the plant physiology, metabolites, ion, and organic solute variation in different plant organs (leaf and/or root) (Barrett-Lennard and Shabala, 2013; Behr et al., 2017).

Developing highly reliable molecular pattern models in plants subject to multiple stresses is a fascinating problem. The transcriptional post-mortem of *Brachypodium dystachion* under the combination of various stresses (drought, salinity, and heat) displayed specific signatures and functional mechanisms in each gene subset (Shaar-Moshe et al., 2017). Transcriptome analysis of combined stress in various plant species revealed a mosaic of transcript behavior. Indeed, these transcriptomic profiles cannot be predicted from the related single stressor (Prasch and Sonnewald, 2013; Coolen et al., 2016; Cohen et al., 2021; Zandalinas et al., 2021). Stress combination raised massively the amount of differentially expressed and regulated genes (DEGs) that are reflected by drastic changes in plant genome and morphology (Desaint et al., 2021). Transcriptome analysis combined with biochemistry showed a significant synergistic response in hexoses biosynthesis following the type of stress combination (Prasch and Sonnewald, 2013). The stress-induced starch degradation leads to an up-regulation of defense gene expression through sugars that are well-known to be signaling molecules (Thalmann and Santelia, 2017). The integration of multiple signals in plants under stress combination that ipso facto leads to a systemic response is far from being taken as an “ipsé dixit”. Heat stress associated with high light in *Arabidopsis* generated systemic signals linked with reactive oxygen species (ROS) that resulted in a systemic response (Zandalinas et al., 2020). The results also suggested that hormonal, metabolomic, and physiological variations affect the integration of systemic signals in *Arabidopsis* under combined stress. The expression of genes encoding jasmonate-responsive transcriptional regulators and those related to ROS-scavengers was strongly down-regulated in JA-deficient mutant aos (*Arabidopsis*) that suggests a prominent role of plant hormone in different specific transcriptional response during combined stress (Balfagón et al., 2019).

Most of the studies cited above are related to the model plant *Arabidopsis* and to the combination of ST, heat stress, high light, or drought. The genetic manipulation of *Arabidopsis* and its use in a greenhouse or experimental chamber is more convenient compared to woody plants for example. Molecular (transcriptomic analysis) responses of plant species under combined hypoxia and ST are missing significantly. The earlier study focused on the tolerance of different wetland tree species has shown the crucial role of ion uptake regulation under SWL (Carter et al., 2006). However, comprehensive transcriptomic integrated with metabolic and physiology analysis in tree specific-organs is needed to understand how trees face combined stress. Here, we investigated the metabolomic and transcriptional patterns in a non-halophyte forest species *Dalbergia odorifera* T. Chen. *D. odorifera* possessed certain waterlogging and salinity tolerance and could be used as preferred tree species in upper parts of the intertidal zones periodically subjected to combined waterlogging and salt stresses. To the best of our knowledge, this is the first study investigating molecular responses (transcriptomic integrated with metabolomic) and patterns of plants under hypoxia combined to salinity. Six woody plant species involving *D*. *odorifera* have been suggested to be suitable for the ecological restoration of flooded watershed areas (Ma et al., 2019). *D. odorifera* exhibited new leaf growth and high fresh biomass growth rate under long-term hydro-culture. It is belonging to the Fabaceae family with a high concentration of phenolics in its organs possessing impressive antioxidant activity. Most of the restoration of flood-related ecosystem projects aimed to increase biodiversity by adding exotic woody plants, and recent initiatives include climate-change adaptation (Nilsson et al., 2018). Indeed, it has been well-established that tree species selected based on tolerance to combined stress are more likely to perform better in stressful areas than those tolerant to a single abiotic stress (Marcar et al., 2002). Here, we reveal the unique metabolomic, transcriptomic, physiological, and hormonal responses of a tree species *D. odorifera* to a combination of hypoxia and saline conditions. Although *D. odorifera* possesses a certain salt tolerance and can grow well under hypoxia, we hypothesize a massive shutdown by SWL of photosynthesis, starch formation, and biomass accumulation. Moreover, it is plausible to expect the same specific pattern mode between the metabolome and transcriptome profiles in *D. odorifera* seedlings under SWL. And since most of the earlier studies focused on combined stress were associated with plant acclimation, shock stress (SWL) applied in the present study would provide significant insights into plant survival and adaptation responses under combined stress.

## Materials and methods

### Plant material and experimental set-up

*D. odorifera* is endemic species to Hainan with significant medicinal value. Two-year-old saplings of *D. odorifera* (500 seedlings) were purchased from a wholesale plant nursery located in Ledong County (18° 42′ 57.91′′ N, 108° 52′ 18.65′′ E) Hainan Province in China. The seedlings grown in a plastic bag were moved into plastic pots filled with red soil, sand and coconut coir (2:2:1, v/v/v). Physicochemical analysis showed that the red soil contained 5.34 mg kg^−1^ of ammoniacal nitrogen, 11.78 mg kg^−1^ of available phosphorus, and 81.72 mg kg^−1^ of available potassium. The seedlings were cut off at a distance of 10 cm from the root; the pots were transferred in a shaded environment in a greenhouse located at Hainan University (20° 03′ 22.80′′ N, 110° 19′ 10.20′′ E). After the young leaves appeared about 5 cm in height, seedlings were placed in normal light conditions and watered every day for two months. A total of 320 healthy seedlings with approximately the same size were divided into two experimental groups. The first experiment was set up with 200 samples in a completely randomized trial in which different stressors were tested for 14 days. The trial comprised eight treatments (25 seedlings each treatment) including a control group (well-watered conditions; CT). The salt treatment was tested at 3 levels; ST1 (100 mM of NaCl), ST2 (150 mM of NaCl), and ST3 (200 mM of NaCl). ST treatments were first imposed at 100% of field capacity and then 100 mL every 2 days during the rest of the trial. For the WL treatment, pots containing seedlings were partially submerged with aqueous solution in a 10 L plastic bucket, and then the water level was maintained 10 cm above the soil surface. SWL treatment was composed by SWL1 (ST1 + WL), SWL2 (ST2 + WL) and SWL3 (ST3 + WL). The pots were placed in the same plastic bucket as WL, and 50 mM NaCl solution was added to replace the evaporated solution. The water potential in 3 leaves sampled from each treatment was measured 1 hour after the stressors were applied, and for 7 days using a Dewpoint PotentiaMeter WP4 (Gene Company Ltd, USA), and values have been reported in Suppl. Table 1a. After 2 weeks of treatments the survival rate was determined and showed in Suppl. Table 1b. Experiment 2 has been set up based on the results of the first experiment. Indeed, the sampling time point has been chosen before the debut of visual stress symptoms and at a high DWP to attempt to glimpse the responses caused specifically by the combination of hypoxia and saline conditions. Moreover, the most suitable combinatory mode of SWL was also selected based on trial 1. Therefore, 4 treatments (30 replicates each) including CT, WL, ST3, and SWL1 were performed during the second experiment and healthy fresh samples were harvested on day 6.

### Morphological and Physiological Characterizations

Measurements of morphological traits were performed on day 6. The biomass accumulation (roots and leaves), leaf surface indexes, and water status were determined in five representative replicates (n=5). Mature leaves at the top were used to measure leaf area, length, and width with a portable area meter LI-3000C (Li-COR, Lincoln, USA). The leaflet water content (LWC) was determined by the following formula: WC (%) = (FW − DW) / FW ∗100, fresh leaflet was weighed (FW) and then dried at 80 °C for 48h. The dried material was measured and recorded (DW). Cell death in root was analyzed spectrophotometrically with the method described by (Baker & Mock, 1994) with Evans blue solution.

Photosynthetic parameters were measured from 9:00 to 12:00 with LI-COR 6400 portable photosynthesis system (LI-COR). The phytohormones such as abscisic acid (ABA), auxin (IAA), and gibberellin (GA_3_) were determined as described in previous research work (Cisse et al., 2022). The non-structural carbohydrates (sucrose, fructose, and glucose), starch, and proline were measured as described by (Pu et al., 2021). The oxidative stress-related enzymes (peroxidases, superoxide dismutase, phenylalanine ammonia-lyase, polyphenol oxidase, and glutathione peroxidase) and molecules (flavonoids, phenols, ascorbic acid, reduced glutathione, and alternative oxidase proteins), and hydrogen peroxide (H_2_O_2_) were determined according to colorimetric Solarbio assay kits. The details related to each assay kit have been provided in Suppl. Table 2 and the superoxide anions was measured as described in (Cisse et al., 2022).

Hierarchical clustering and silhouette analysis were used to determine the distance between the resulting clusters from different morpho-physiological parameters, and according to their plant-related function. Thus evaluating which stress between ST and WL affects more *D. odorifera* seedlings under SWL. The hierarchical clustering was based on Euclidean distance, and then Ward’s linkage method was used to minimize the increase in the error sum of squares. The silhouette plot with a range between (−1, 1) showed how close each group of a parameter (n = 5) belongs to its own cluster (group of data from one treatment) and to another cluster. If the coefficient is close to +1 then the cluster is afield from the other clusters belonging to the rest of the treatments. A null coefficient implies that the cluster is very close to another neighbor cluster, and a negative coefficient indicates that those data from one treatment belong to another cluster.

### Safranin Fast Green (SFG) and Transmission Electron Microscope (TEM) staining

Leaflets including major veins and roots (n=6) collected on day 6 were cut and fixed in 50 % FAA fixative solution for 24h. The samples were deparaffinized with pure BioDewax and Clear Solution for 20 min, and with pure ethanol for 5 min, then finally with 75% ethanol. The slides were stained in fast green dye solution for 1-5 minutes, washed, and soaked in 1% hydrochloric acid and alcohol for 10s. Thereafter, slides were stained in saffron dye solution for 1-5s and then put into 100% ethanol for dehydration. Finally, the slides were immersed in xylene for 5min, sealing with neutral resin. Samples were observed under orthostatic microscope (Nikon Eclipse E100, Nikon, JAPAN) and the images were analyzed with CaseViewer software (3DHISTECH Ltd., Budapest, HUNGARY).

Fresh leaf tissues of *D. odorifera* under different stress treatments were analyzed by TEM to visualize the variations of the chloroplast, starch granule, and plastoglobuli. Briefly, a sharp blade was use to harvest 1 mm3 of sample and transferred into an EP tube with fresh TEM fixative (Servicebio, G1102) for fixation. The samplings were placed at room temperature for 2h and then fixed at 4°C. Thereafter, the tissues were washed using 0.1 M phosphate buffer (PB) (pH 7.4) 3 times (15 min each). About 1% of osmium tetroxide (OsO4) in 0.1 M PB (pH 7.4) was used in post-fixation for 7 h at room temperature, and then the OsO4 was removed with 0.1 M PB (pH 7.4) for 3 times (15 min each). The samples fixed in OsO4 solution were dehydrated in graded ethanol and mixed ethanol: acetone successively. The ethanol series comprised 30%, 50%, 70%, 80%, 95%, 100%, and 100% of ethanol for 1h each. And then with ethanol: acetone (3:1, 0.5h), ethanol: acetone (1:1, 0.5h), ethanol: acetone (1:3, 0.5h), and pure acetone for 1h. The resin penetration and embedding were performed with pure EMBed 812 (SPI, 90529-77-4) and a series of mixed solution (acetone and EMBed 812) as following: acetone: EMBed 812 (3:1) for 2-4 h at 37°C, acetone: EMBed 812 (1:1) overnight at 37°C, acetone: EMBed 812 (1:3) for 2-4 h at 37°C and 100% of EMBed 812 for 5-8 h at 37°C. The embedding samplings were transferred into a 65°C oven (48h) for polymerization, and then the blocks were cut to 60-80nm thin with an ultramicrotome (Leica UC7). The tissues were fished out onto the 150 meshes cup rum grids with formvar film. For the staining 2% of uranium acetate (avoid light staining) saturated alcohol solution was used for 8 min, and then the cup rum grids was rinsed with 70% ethanol for 3 times and with ultra pure water for 3 times. At last 2.6% lead citrate (avoid CO2 staining) was used for 8 min, and then samples were rinsed with ultra pure water for 3 times. After dried, the cup rum grids are observed and capture with TEM (HITACHI, HT7800/HT7700).

### Widely targeted metabolomics assay

Freeze-dried leaf samples were crushed into powder using a mixer mill (MM 400, Retsch) with a zirconia bead for 1.5 min at 30 Hz. About 100 mg of powder was weighed and extracted at 4 °C with 1.0 mL 70% aqueous methanol and then centrifuged at 10, 000 g for 10 min. The sample extracts were absorbed (CNWBOND Carbon-GCB SPE Cartridge, 250 mg, 3 mL; ANPEL, Shanghai, China) and filtrated (SCAA-104, 0.22μm pore size; ANPEL) before LC-MS analysis. LC-ESI-MS/MS system was performed to analyze the extracts and the analytical conditions were as follows, HPLC column, Waters ACQUITY UPLC HSS T3 C18 (1.8 µm, 2.1 mm*100 mm); solvent system, water (0.04% acetic acid): acetonitrile (0.04% acetic acid); gradient program, 95:5V/V at 0 min, 5:95V/V at 11.0 min, 5:95V/V at 12.0 min, 95:5V/V at 12.1 min, 95:5V/V at 15.0 min; flow rate, 0.40 ml/min; temperature, 40°C; injection volume: 2 μl. The effluent was connected to an ESI-triple quadrupole-linear ion trap (Q TRAP)-MS. Triple quadrupole (QQQ) scans and LIT were acquired on a Q TRAP mass spectrometer, API 6500 Q TRAP LC/MS/MS System. The system was equipped with ESI Turbo Ion-Spray interface and functioning in a positive ion mode and managed by Analyst 1.6.3 software (AB Sciex). The electrospray ionization (ESI) parameters were set as follows: the collision gas (N2) was set 5 psi; ion spray voltage 5500 V; source temperature, 500 °C; ion source gas I, gas II and curtain gas were set at 55, 60, and 25.0 psi, respectively. QQQ scans as multiple reaction monitoring (MRM) experiments with nitrogen as collision gas were required. The de-clustering potential (DP) and collision energy (CE) for individual MRM transitions was performed with further DP and CE optimization. For each period according to the metabolites eluted within this period, a specific set of MRM transitions were monitored. The secondary spectral data obtained were qualitatively analyzed based on public metabolite database (MassBank, KNApSAcK, Metlin, MoTo DB and hmdb) and a self-built database MetWare database (MWDB). The annotation of differential metabolites and metabolite enrichment pathway analysis were performed with Kyoto Encyclopedia of Genes and Genomes (KEGG) database (http://www.genome.ad.jp/kegg/)

### Transcriptomic analyses

We performed RNA-seq to dissect the global transcriptional adaptations in *D*. *odorifera* leaflet under single and combined stresses. The transcriptomic assay involving RNA extraction and sequencing, data processing, and analysis was performed in four replicates (fresh mature leaves from the top). Briefly, fresh leaf samples were harvested on day six from four replicates and immediately frozen in liquid nitrogen, and the total RNA was extracted using the TRIzo Kit (Promega, Beijing, China). Sample contamination and RNA purity were assessed using NanoPhotometer (Thermo Fisher Scientific) at optical densities 260/280 and 260/230. The RNA integrity was detected accurately with Agilent 2100 Bioanalyzer (Agilent Technologies, CA, USA). Poly (A) mRNA was isolated from total RNA samples with Magnetic Oligo (dT) beads and used for mRNA-sequencing library construction. To select cDNA fragments (preferentially 250–300 bp in length), library fragments were purified with an AMPure XP system (Beckman Coulter, Beverly, MA, USA). The usability of the RNA-Seq data was verified through quantitative real-reverse transcription PCR (qRT-PCR). The mRNA-sequencing libraries were sequenced using Illumina HiSeqTM 2500 sequencing platform (Illumina Inc., San Diego, USA). The data obtained from the sequencer were converted into sequence data (raw reads) by CASAVA and stored in FastQ files format. Sequencing data was filtered using an ultra-fast all-in-one FASTQ preprocessor (fastp) according to (Chen et al., 2018), in order to ensure the reliability of bioinformatics analysis and to obtain high-quality data for subsequent analysis.

The Base quality assessment of each sample was inspected using FastQC (http://www.bioinformatics.babraham.ac.uk/projects/fastqc). Trinity was used to assemble the filtered high-quality sequencing data to obtain the transcriptome, which is used as a reference sequence for subsequent differential expression analysis. The unigene was extracted from the transcriptome, and the relevant information of the transcript and unigene were counted, as shown in Suppl. Table 3a. CoDing Sequence (CDS) prediction was performed before the gene function annotation. Six databases such as KEGG, GO, NR, Swiss-Prot, trEMBL, and KOG were used to annotate the predicted protein sequences. The conditions used for BLAST comparison database are Evalue le-5 below, identity above 30%, and sequence coverage above 30%. The software package RSEM (RNA-Seq by Expectation Maximization) was set up to quantify gene and isoform abundances. Thus, RSEM software was used to quantify the fragments per kilobase of transcripts per million fragments mapped (FPKM) of each corresponding gene. Differential expression genes (DEGs) were detected based on the fold change (FC) of the FPKM values. The false discovery rate (FDR) was fixed at ≤5% and used to define the p-value threshold (Love et al., 2014). The threshold for screening DEGs was arranged at an absolute value of log2FC and employed to define the p-value threshold (Benjamini & Hochberg, 1995). The threshold foclusterProfiler R package, GO, and KEGG pathway enrichment analyses of the DEGs were performed.

### Expression Patterns of genes and metabolites

The characterization of the shared metabolites and gene patterns will significantly contribute to our understanding of the uniqueness of plant response under combined stress compared to a single stressor. The accelerating availability of data that described molecular pathways involved in plant response under single stress allowed us to better understand the function of various specific genes, proteins, or tissues in plants following the conditions in their natural habitat. However, plants in their natural environment confront various simultaneous environmental conditions. Few research studies such as in (Shaar-Moshe et al., 2017) attempted to describe the common transcripts pattern mode in plants under combined stress. Here, we provided a global molecular pattern mode including metabolome and transcriptome in a tree species under combined stress. An Interactive Venn diagram was used to identify common differential genes and metabolites expression. First, differential metabolites were screened at fold change ≥ 2 and fold change ≤ 0.5, if the difference between the control and the stress group is more than 2 in up-regulated metabolites or less than 0.5 in those down-regulated, the difference between the metabolites is considered significant. After that, based on the description above, selected metabolites with (variable importance in projection) VIP≥1 were chosen. The common transcript expression patterns were determined based on log2FC and FDR according to (Shaar-Moshe et al., 2017). We classified each pattern found in the transcriptome and metabolome as synergistic, additive, dominant, neutral, minor, unilateral, antagonistic and non-assigned. The standard deviation SD was set up at 0.50 based on the SD among the unique differential transcripts and metabolites in SWL. Metabolites or transcripts assigned to synergistic, additive and dominant modes were either up- or down-regulated among the single and combined stresses. The sum of the individual-stressor metabolite and transcript was calculated, and then the pattern mode in SWL is synergistic if SD between the Sum is > 0.50 otherwise its additive mode. Based on the SD if the metabolite or transcript log2FC of one of the single stress is equal to SWL then the pattern is dominant. The neutral pattern mode corresponds to an SD ≤ 0.50 among the log2FC of each single stress group compared to SWL, whether they were either up- or down-regulated. The minor combined-stress modes were determined based on a lower (±SD) log2FC than one or both single stress. Metabolites and transcripts with opposite expression patterns were assigned to unilateral or antagonistic modes. The mode is antagonistic if both expression patterns in single stress are opposite to SWL (±SD). Unilateral mode included metabolites or transcripts from SWL that were only followed an opposite expression patterns to one of the single stress (±SD).

### Statistical Analyses

Analyst 1.6.3. Multivariate statistical analysis methods was used for mass spectral data processing, including principal component analysis (PCA) and orthogonal partial least squares discriminant analysis (OPLS-DA) according to the method describe by (Eriksson et al., 2006). The data from metabolite analysis were normalized and R software (www.r-project.org/) was used to perform cluster analysis (Hierarchical cluster analysis, HCA). The data belonging to the physiological and morphological indexes were statistically differentiated with two-way ANOVA and Tukey’s honestly significant difference test (Graph Pad prism 9.0.0). Data were expressed as means ± SD (5 replicates), and significant differences between means were determined at a p-value ≤ 0.05.

## Results

### Global metabolome multivariate statistical analysis showed how combined stress was far from the control group but closer to one of the single stress

The visualization of complex molecular data can be simplified and dimensionally reduced based on multivariate statistical analysis to summarize the different characteristics of the metabolome profile and determine their pattern. This step is crucial to define how the metabolites clusters are organized and close to each other in plants under single and combined stress. The cluster analysis (Fig. 1A_1_, A_2_) showed the metabolites accumulation and variation in the *D. odorifera* leaflet. Results represented in heat maps showed that the metabolites in SWL had similar trends compared to ST. Based on Pearson’s correlation coefficient (r2 and r1) SWL displayed a range of 0.98 to 0.96 correlation with CT or WL, compared with the correlation between SWL and ST with a range of 0.98 to 1, suggesting that metabolites variation under combined stress was led by ST (Fig. 1A_1_). The highest homogeneity within the same cluster or compared to another cluster was found in the ST group while WL and SWL clusters showed the highest heterogeneity. The principal component analysis (PCA) reflected that a major difference existed between metabolite levels within control, single, and combined stress treatments (Fig. 1A_3_, A_4_). The first component (PC1) separated CT and SWL while the second component (PC2) separated CT and WL. Interestingly, the ST group had at least one outliner regarding both principal components. Indeed, among the 3 groups from ST, the PC1 separated CT and two groups of ST combined with 3 groups of SWL (Fig. 1A_3_). To verify the consistency of the metabolome profile clustering the data was subjected to a grouped principal component analysis 3D plot (Fig. 1A_4_). The result showed that PC1 split the SWL group from the rest of the treatments. However, the PC3 separated two groups from WL and SWL into two groups CT and ST. Based on the 3D PCA plot only the separation of SWL from the rest of the treatment was consistent with the results shown in the 2D PCA plot. The UPLC-MS/MS detected 572 metabolites in each treatment, and then a total of 137 differential metabolites (DMs) were screened in SWL following a fold change (FC) ≥ 2 or FC ≤ 0.5, and VIP ≥ 1. Intersections among down- (Fig 1B_1_) and up-regulated (Fig 1B_2_) differentially expressed metabolites across single and combined stress showed that there were more unique DMs (27 down- and 31 up-regulated) compared to common metabolites regulated under ST, WL, and SWL. The combined stress resulted in an increase of DMs compared to those in single stress (Fig 1B_3_, Suppl. Table 4, 5). The interactive Venn Diagram (Fig 1B_4_) of DMs between single and combined treatments showed less variation between SWL and ST than the other groups. The common shared down-regulated DMs was composed of different class such as flavone, amino acid and derivatives, and organic acids and derivatives (Suppl. Table 4) while the common up-regulated DMs comprised the carbohydrates, flavonoid, alcohols, phenylpropanoids, proanthocyanidins, flavone, organic acids and derivatives, isoflavone, alkaloids, nucleotide and derivates, and lipids (Suppl. Table 5). The unique regulated DMs under combined stress outnumber those in ST or WL in terms of diversity of metabolites compounds and family classes. Combined stress-induced lipids down-regulation such as MAG (18:3) isomer5, MGMG (18:2) isomer2, DGMG (18:1), 14,15-Dehydrocre-penynic acid, 9-Hydroxy-(10E,12Z,15Z)-octadecatrienoic acid or 13-HOTrE (Suppl. Table 4) which are crucial to plant photosynthesis inhibition. Such phenomena were not explained by the metabolomic profile in *D. odorifera* leaflet under single stress. These lipid compounds were up-regulated in both stresses applied individually (Suppl. Table 5), which showed a specificity of regulation due to combined stress. The unique shared up-regulated DMs under ST (5) were massively lower compared to those in SWL (31) or WL (22). To further understand the synergistic effects on the metabolome profile caused by combined stress, a quantitative analysis based on log2FC (Suppl. Fig 1A_1-3_) and VIP scores (Suppl. Fig 1B_1-3_) of the detected metabolites has been conducted. The comparison between SWL and CT, ST, or WL showed the uniqueness of the metabolome profile in *D. odorifera* leaflet under combined stress.

**Figure 1.**
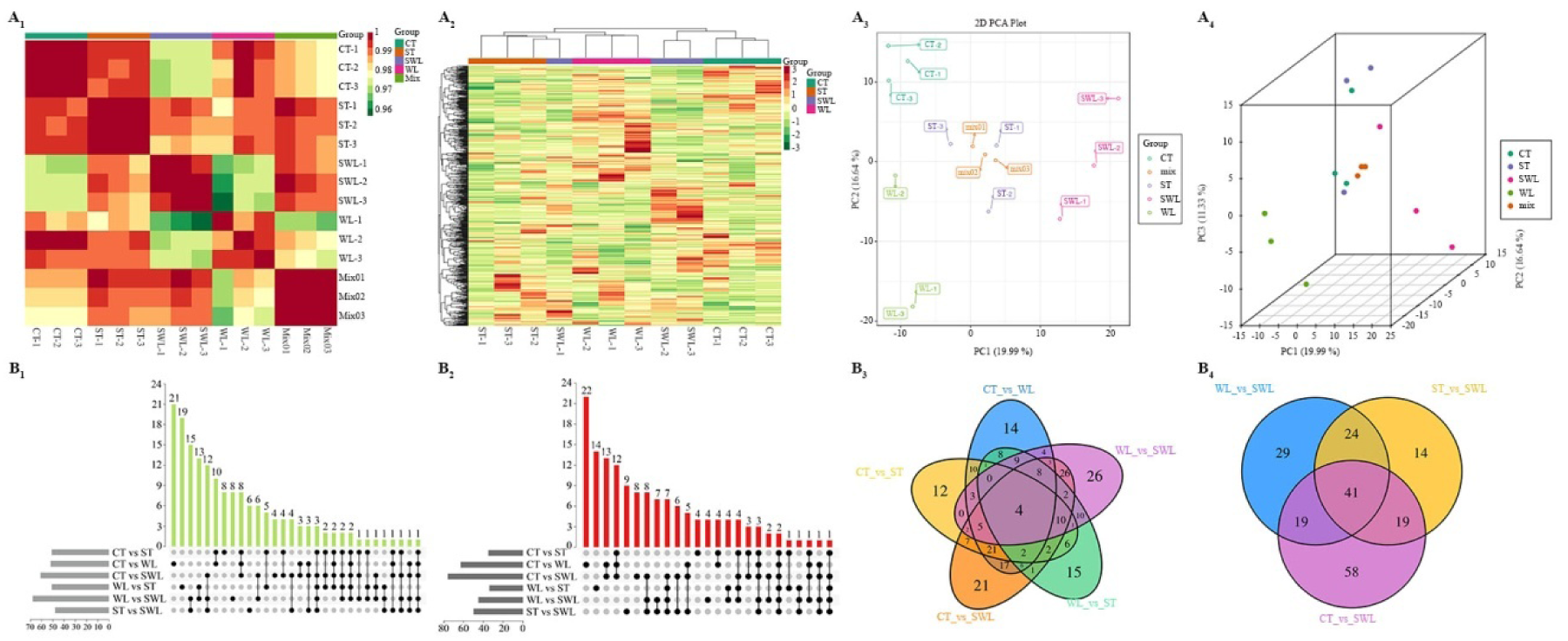
Overview of the metabolomic data assessment in response to single and combined stress, (**A_1_**) Correlation matrix of the whole metabolome data set using three biological replicates and Pearson’s distance as the metric. (**A_2_**) Metabolomic of all samples clustering heatmap, various colours are obtained after normalization (red means high content, green means low content). (**A_3_**) 2D plot of the principal component analysis (PCA) of the metabolites data set regrouping all treatments and (**A_4_**) 3D plot. Intersections among down- (**B_1_**) and up-regulated (**B_2_**) differentially expressed metabolites across single and combined stresses. Number above bars showed the number of metabolites within each intersection. Venn diagrams display the relationship among different metabolites in various comparative groups, including one with CT and WL as a control (**B_3_**) and a second with SWL as a control (**B_4_**). Treatments are presented as follows: control (CT), salinity (ST), waterlogging (WL), and combination of WL and ST (SWL).

### Combined stress resulted in a massive impact on the plant leaflets’ transcriptome

RNA-seq has been used to dissect how combined stress compared to single stress affects the transcriptome in *D. odorifera* leaflet. The transcript length distribution comprised a total of 703596 transcripts and 419093 Unigene. A heat map of the Pearson correlation coefficient was determined between all sampling and was displayed in Fig. 2A_1_. The relationship between the transcripts expressed for a given gene in the combined treatment versus CT, ST, or WL was smaller than that shown among the single treatment (ST or WL) versus CT. Indeed the minimum coefficient found in SWL versus ST or WL was 0.30 and 0.34, respectively. It showed a consistent minimum coefficient between CT and SWL, but a comparison among combined and single stress showed at some point a strong correlation with a maximum coefficient number of 0.93 in the ST group and 0.90 in the WL samples. The density distribution plot (Fig. 2A_2_) and side-by-side box plots (Fig. 2A_3_) showed a minimal difference in the all-around transcripts of dissimilar samplings, and most of them had log10 (FPKM) values between −2 and 2. Both single stress samples were distributed between SWL and CT; at this stage it was impossible to detect a cleaner separation among the four treatment groups. It was also difficult to evaluate how closely the transcript clusters from both single stresses were close to the combined stress. To further understand the pattern between single and combined stress, differential gene expression analysis (DESeq2) has been performed. The results showed an impressive number of transcripts regulated that were involved in SWL (14053) compared to WL (4111) or ST (2670). In all three treatments compared to CT, the down-regulated DEGs outnumbered the up-regulated transcripts (Suppl. Table 3c). Hierarchical clustering of the most highly and low expressed genes determined with RNA-seq in *D. odorifera* leaflets samples showed that the pattern of clusters from biological replicates of ST, WL or CT compared to combined stress are different (Fig. 2B_1-3_). The outcomes revealed that transcripts with identical or similar expression patterns under different treatments were clustered into classes, and these similar transcripts could be regrouped for further analysis. The overlapping area of the Venn diagrams (Fig. 2C_1-3_) of the differential genes resulted in different groups that indicated the intersection between combined stress and the other treatments. This step is crucial to isolate the commonly shared genes among WL, ST, and SWL. For instance, if the reference of DEGs comparison is CT, it was noticed 519 common genes between ST, WL, and SWL (Fig. 2C_2_). Volcano plots are a helpful representation to visualize the results of differential expression analyses (Fig. 2A, C) as well as the MA-plots showed in Fig. 2B, D. In the volcano plots, the abundance of down- (green) or up-regulated DEGs (red) and non DEGs (black) were compared across all treatments based on the (log 2 Fold Change) log2FC for each gene, along with the false detection rate (FDR < 0.05) that represents the statistical significance of the differential expression test. The MA plots showed the log2FC attributable to a given variable over the mean of normalized counts for all the samplings from the RNA-seq data set. Figures A_3_ and B_3_ symbolized the synergistic effects of combined stress in plants compared to a single stress. Indeed, whether the DEGs were either up- or down-regulated, their abundance under combined stress was massively impressive. The high presence of regulated DEGs (red color point) in combined stress (Fig. 3B_3_) showed their prominent role in stress tolerance compared to those in single stress (Fig. 3B_1-2_). To determine the pattern of the regulated DEGs in SWL regarding its individual component, we performed volcano (Fig. 3C) and MA (Fig. 3D) plots among WL vs ST, ST vs SWL, and WL vs SWL. The results showed an abundance of significant DEGs in the group of WL vs SWL compared to the other groups. Thus the genes regulated in SWL were closer to those in ST compared to WL.

**Figure 2.**
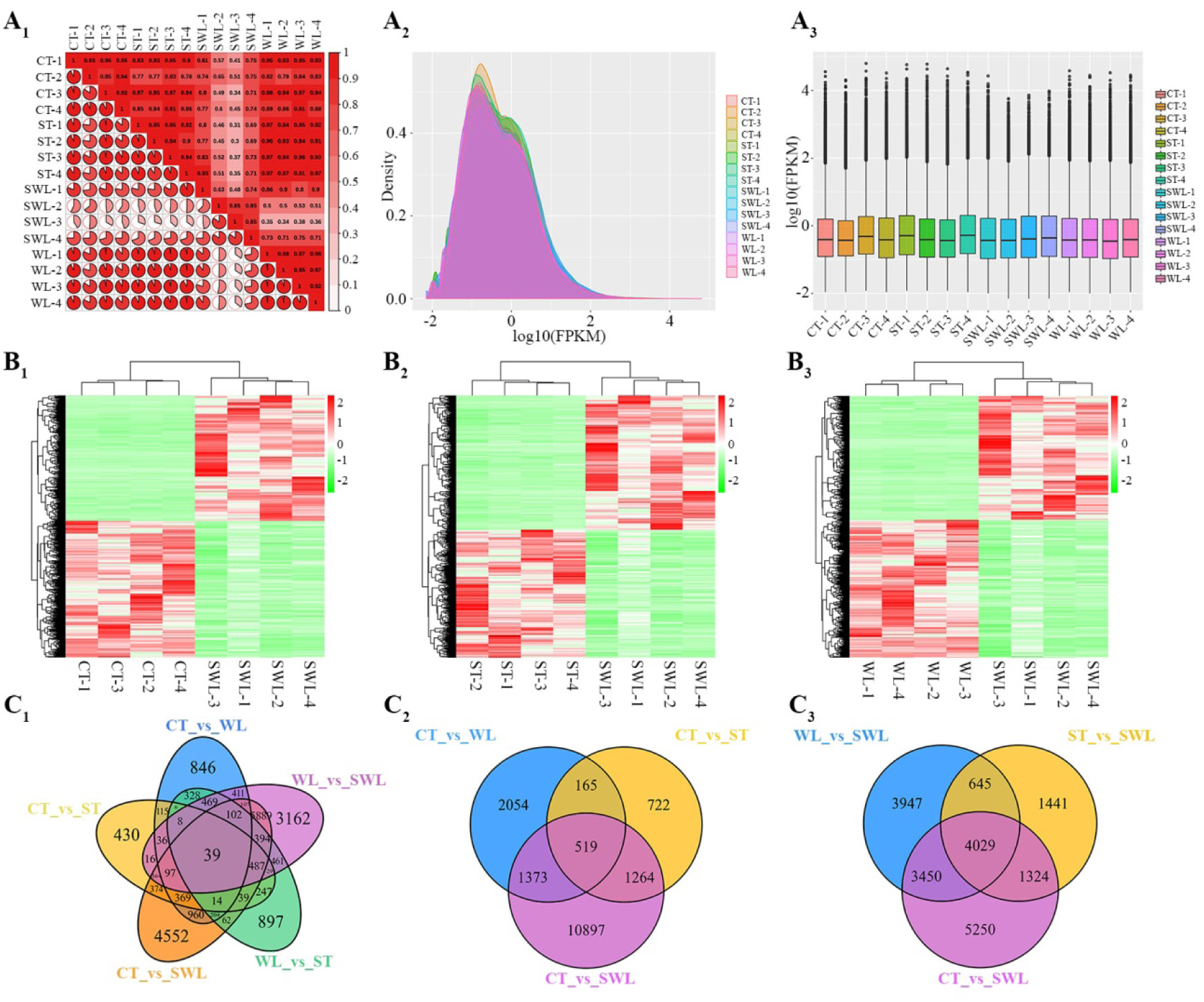
Summary of the transcriptomic data assessment in response to single and combined stress, Pearson’s correlation diagram regrouping all treatment from different treatments (The closer the correlation coefficient is to 1, the higher the similarity of expression patterns between samples.) and according to the gene expression level FPKM, the correlation coefficient of samples within and between groups was calculated and drawn as a heat map (**A_1_**). Gene expression density map regrouping all samples based on log10FPKM (**A_2_**) and box plot of gene expression distribution map (**A_3_**). Cluster heat map analysis of expressed genes among CT vs SWL (**B_1_**) ST vs SWL (**B_2_**) and WL vs SWL (**B_3_**). Interactive Venn diagram of differential gene results for different groups, overlapping areas indicate intersections between various combinatory groups, and numbers represent the number of differential genes (**C_1_**, **C_2_** and **C_3_**). Treatments are presented as follows: control (CT), salinity (ST), waterlogging (WL), and combination of WL and ST (SWL).

**Figure 3.**
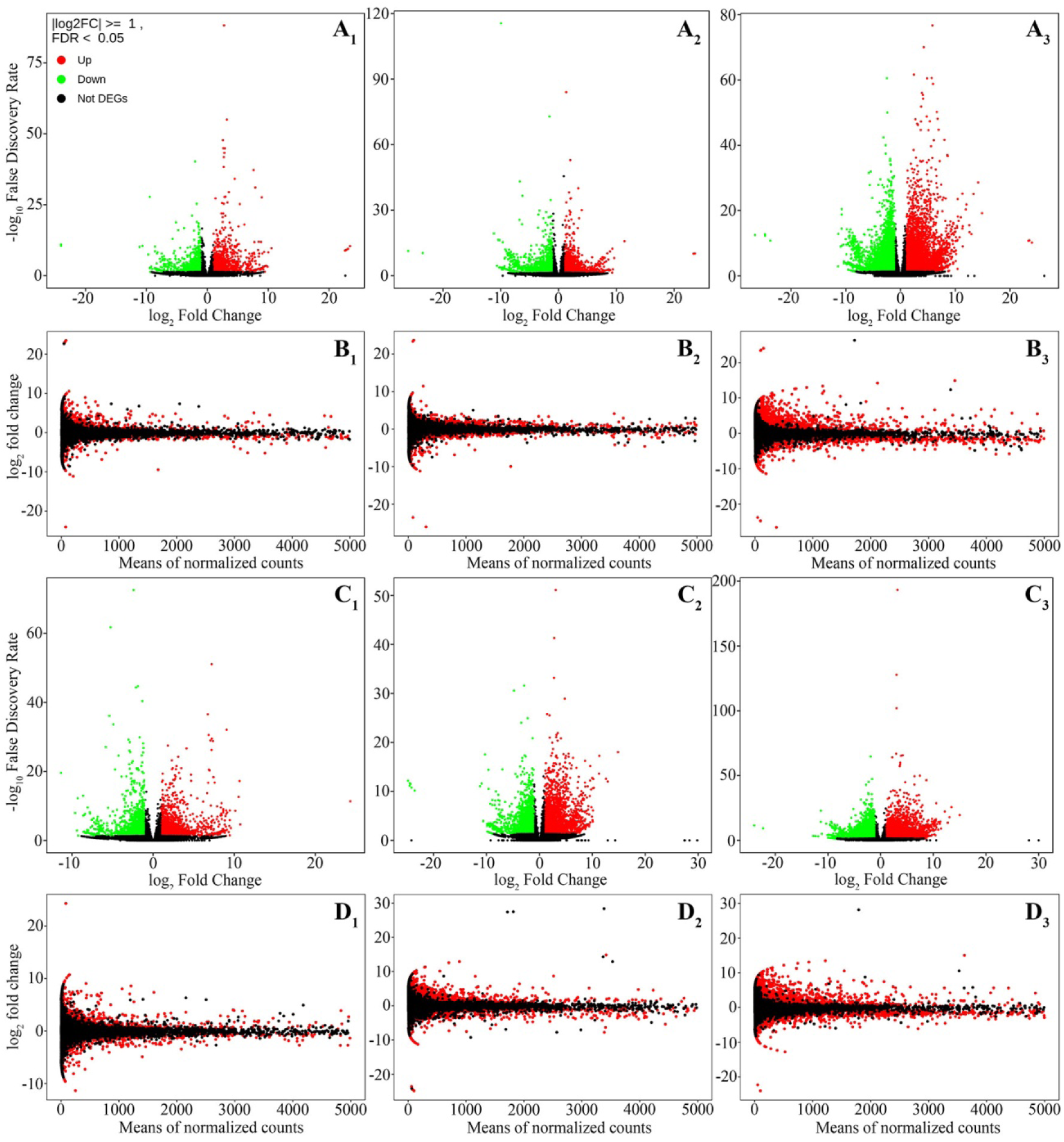
Volcano plots of DEGs from ST (**A_1_**), WL (**A_2_**) and SWL (**A_3_**) treatments compared to CT based on the log2FC and the false detection rate (FDR < 0.05), the green colour point corresponded to the down-regulated genes and red colour those up-regulated. MA-plots of DEGs from ST (**B_1_**), WL (**B_2_**) and SWL (**B_3_**) treatments compared to CT, transcripts will be coloured in red if the FDR is less than 0.05 and the rest of DEGs were coloured in black. Volcano plots of DEGs from SWL compared to WL (**C_1_**) and ST (**C_2_**) and from STB compared to WL (**C_3_**). MA-plots of DEGs from SWL compared to WL (**D_1_**) and ST (**D_2_**) and from ST compared to WL (**D_3_**). Treatments are presented as follows: control (CT), salinity (ST), waterlogging (WL), and combination of WL and ST (SWL).

### Integration between the core biological functions of genes and synergistic pattern of *D. odorifera* leaflet under combined stress

The previous results showed that combined stress raises a tremendous amount of transcripts compared to a single stress. The mobilization herculean of these genes under combined stress is a fundamental response in plants for combined-stress tolerance and survival. And to qualitatively determine the role of each transcript and provide more insights into their pattern particularly under combined stress, Gene Ontology (GO), Kyoto Encyclopedia of Genes and Genomes (KEGG), and Eukaryotic Orthologous Groups (KOG) databases have been used to classify and annotate them. Functional gene categories of antioxidants (phenolic compounds and proline), photosynthesis, carbohydrate metabolism, DNA and RNA processes, senescence, and necrosis, signaling pathways, and transporters were selected based on KEGG classification to analyze their expression pattern in combined stress. The KEGG classification of DEGs in each treatment has been shown in Suppl. Fig 2-4, the pattern of each functional gene category was visualized in a heat map (Suppl. Fig 5-6). Interestingly the sum of the number of DEGs involved in metabolic pathways under single ST (257) and WL (383) was multiplied by two under combined stress (1349). The same trend was observed in the transcripts involved in the biosynthesis of secondary metabolites. *D. odorifera* as a medicinal woody plant with an impressive number of phenolic compounds, showed a massive regulation of functional genes involved in phenylpropanoids, flavone, isoflavone, phenylalanine, or carotenoid pathways Suppl. Fig 5A. These transcripts are crucial in phenolics accumulation that plays a role of antioxidants in plant system defense as well as proline or glutathione. From receipting signals from harsh conditions to the response of plant cells to provide stress tolerance, the results highlighted a quasi-perfect impressive synergistic pattern of each functional gene category involved in plant stress tolerance under combined stress. Indeed, the ABC transporters, MAPK, calcium, NF-kappa B, Toll-like receptor, and NOD-like receptor signaling pathways, as well as plant hormone signal transduction showed a harmonious pattern in combined stress Suppl. Fig 5A. The same patterns were observed in carbon and photosynthetic-related functional genes (Suppl. Fig 6A) and those related to cellular senescence and necrosis or the circadian rhythm – plants (Suppl. Fig 5C). The whole DEGs of each treatment were distributed in three categories of GO (molecular function, cellular component, and biological process) to confirm the consistency of DEGs pattern between combined and single stress found in KEGG enrichment. The different functional gene categories in GO classification (Suppl. Fig. 7-9) showed the same pattern as those in KEGG analysis. However, statistically, the scatter plot of KEGG (Suppl. Fig. 10) and GO enrichment (Suppl. Fig. 11-13) displayed a significant difference in the most enriched pathways between combined and both single stress that had the most contribution. The KOG (Suppl. Fig. 14) as well as GO and KEGG functional annotation and enrichment analysis displayed a massive enrichment under SWL compared to ST or WL. Also, various biological functions were only enriched in SWL. Thus combined stress generates a significant quantitative and qualitative transcriptional response compared to a single stress.

### Characterization of the metabolomic and transcriptional pattern mode in combined stress

One of the central issues in plant stress tolerance nowadays is the understanding of the systemic responses in plants under combined stress by integrating its entire individual component. Although several researchers have attempted to describe the transcriptional changes in plants under combined stress, not many research papers have brought together the individual parts into a global pattern based on each metabolite or gene pattern. The common metabolite and transcriptional pattern in combined stress based on each molecular constituent in their respective single stress and the standard deviation (SD) showed seven pattern modes (Suppl. Table 6-7). Indeed, the similitude in metabolite and gene expression under single and combined stresses and their functions in plant leaflets under SWL was estimated through metabolomic and transcriptional pattern analysis. Fig. 4A showed the SD among Log2FC of metabolite and gene expression in ST, WL, or their sum compared to SWL. The color red (up-regulation) or green (down-regulation) refers to the regulation mode of the metabolites and genes in different treatments. The total of common metabolites detected in all treatments regardless of the VIP and FC was equal to 569, and most of the metabolites in SWL were in a neutral mode (53.78%), as clearly visualized in Fig. 4B. The antagonistic mode in which metabolites were oppositely regulated between single and combined stress represented 7.38% of the metabolites. The number of regulated metabolites in antagonistic mode was higher than those in synergistic added to additive mode (6.15%). Broadly speaking, the well-known synergistic pattern of combined stress in plants does not apply to the regulated metabolites in *D. odorifera* leaflets under SWL. The rest of the assigned metabolites were distributed in the dominant (5.80%), minor (6.68%), and unilateral (11.07%) modes. To find a more informative metabolic pattern in SWL, common DMs were selected based on VIP (VIP≥1) and FC ≥ 2 (up-regulated metabolites) or FC ≤ 0.5 (down-regulated metabolites). A total of 32 DMs were found and the neutral mode contained most of the DMs (50%). The synergistic and additive modes combined together represented 25.01% of the DMs, and the rest were distributed in the minor mode (Fig. 4C). Interestingly, the common DEGs distributed in the synergistic and additive modes (31.03%) had approximately the same percentage as those in the neutral mode (31.98%). These results indicated that many DEGs in SWL compared to ST or WL were massively up or down-regulated (Fig. 4D). The fold changes (FC) that expresses the gene expression level followed a synergistic or additive pattern in *D. odorifera* leaflets under combined stress. There was an important number of DEGs that followed a dominant mode (21.97%) in SWL, and 10.02% of DEGs expression level with the same regulation mode in SWL was significantly lower in SWL compared to ST or WL. The number and profile of common DEGs compared to those in DMs or the overall metabolites were quite different. However, a high number of common regulated DMs or DEGs were expressed similarly in *D. odorifera* leaflets under combined or single stress.

**Figure 4.**
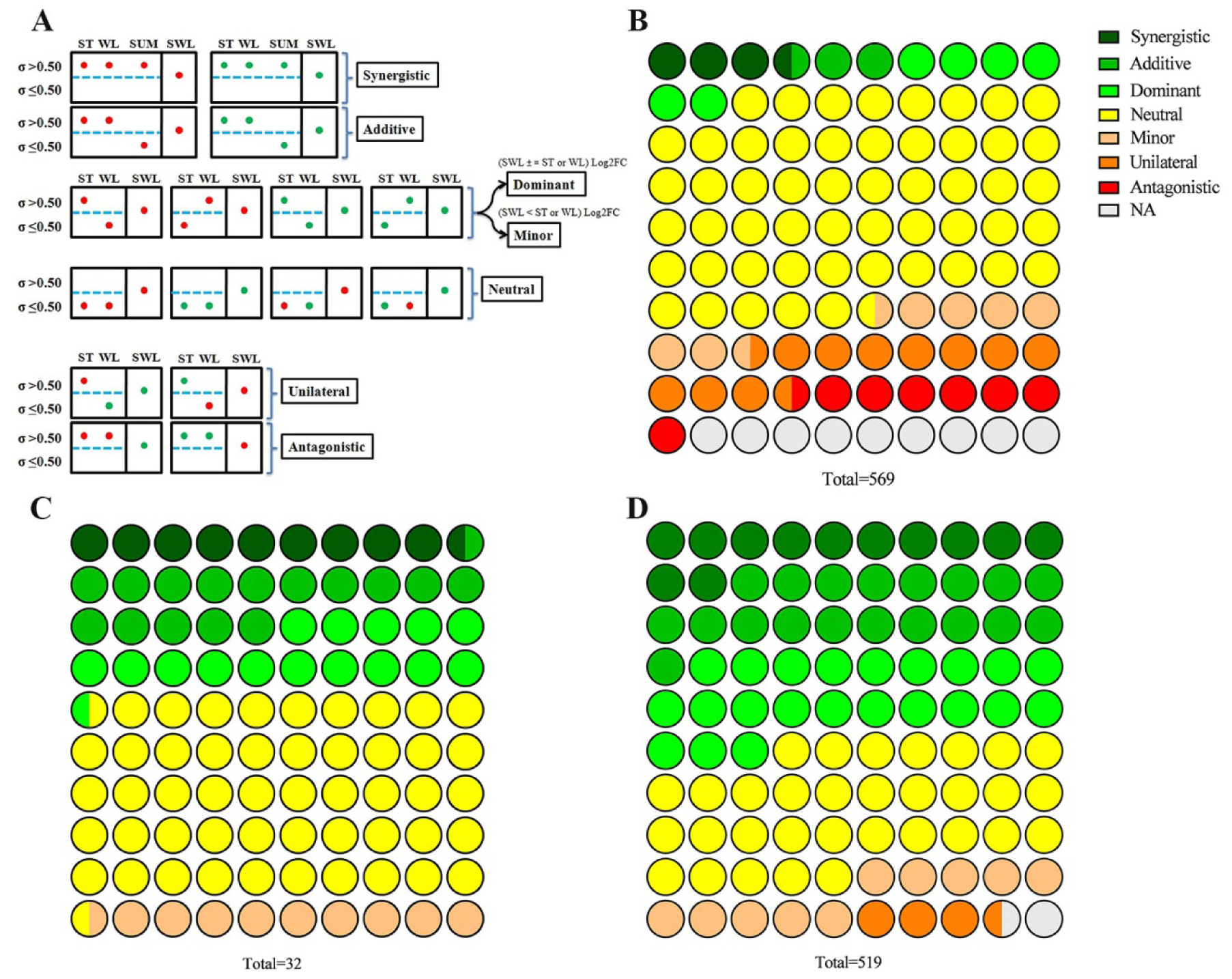
Metabolomic and transcriptional patterns based on log2FC, regulation status (down or up) and standard deviation (SD ≤ 0.50) of *Dalbergia odorifera* responses to single and combined stress. Illustrative schema of molecular expression (metabolites and genes) that showed seven pattern modes (synergistic, additive, dominant, neutral, minor, unilateral, and antagonistic) found in *D. odorifera* under combined stress compared to each single stress (**A**). 10 x 10 dot plot showing the pattern modes; in SWL of the whole metabolites detected (569 metabolites) (**B**) and common differentially expressed metabolites (32) (**C**) based on the comparison in ST and WL, and in the common differentially expressed transcript (519) (**D**). Treatments are presented as follows: control (CT), salinity (ST), waterlogging (WL), and combination of WL and ST (SWL).

### Integration among hierarchical cluster, transcriptional pattern and indexes related to photosynthesis

Euclidean distance metric was determined among the CT, ST, WL, and SWL that comprised the monomers (glucose, sucrose, and fructose) and polymeric carbohydrates (starch), number of starch granules (chloroplast), net photosynthetic rate, stomatal conductance, and plant hormones (ABA, GA_3_, and IAA) indexes. The dendrogram showed two different clusters regrouping the CT in one hand and a second group containing ST, WL, and SWL (Fig. 5A). The silhouette analysis showed the degree of cohesion and separation of the different clusters among different treatments. Fig. 5B showed silhouette values close to 1 in CT and SWL treatments compared to those in ST or WL. The values found in ST and WL clusters depicted two possibilities that included the proximity of CT or/and SWL groups, but a look back to Fig. 5A showed that both clusters were close to SWL. To gain a better understanding of plant leaflets’ photosynthetic variations under combined stress the chloroplast ultra-structure analysis of *D. odorifera* leaflets has been conducted by transmission microscopy (Fig. 5C-D). The density covered by the chloroplasts in the leaflet cells belonging to the CT group was massively more important than those in single or combined stress. The images shown in Fig. 5C-D symbolize the synergistic effects of combined stress in plants by the reduction of starch granule size and number in one hand, and the increase of osmiophilic plastoglobuli (OP) in SWL. Indeed the OP thylakoid-associated lipids that actively participate in thylakoid function from biogenesis to senescence were massively accumulated in SWL-treated leaflets. The synergistic degradation of the polymeric carbohydrates in SWL-treated leaflets (Fig. 5C-F) was correlated with an increase of monomers such as sucrose and glucose (Fig. 5G-H) or fructose (Suppl. Fig. 16A) compared to CT. But glucose and sucrose accumulations in WL or ST were not significantly decreased compared to those in SWL. Thus there was no synergistic effect of SWL in glucose and fructose biosynthesis compared to the sucrose that followed a synergistic accumulation in combined stress regardless of ST or WL. To integrate transcriptional pattern modes into photosynthesis processes under combined stress, GO classification was conducted to regroup several genes and their isoform in six functional groups. The RNA-seq results provided genes and their isoform, and then we determined the frequency (ω) of each gene and its encoded proteins (Fig. 6A-D). The photosynthetic pattern modes have been determined in the common DEGs regrouping either both or one stress compared to SWL. If genes or its isoform were expressed only in SWL a zero was attributed to both rows of single stress, and thus the pattern mode was de facto synergistic. The results in Suppl. Table 7 showed different pattern modes of DEGs related to the photosystem I, photosystem II, photosynthesis light harvesting, plastoglobule biosynthesis, polysaccharide catabolic process and carbohydrate transport. Interestingly, in each functional subset gene, the synergistic effect of combined stress was sovereign (Fig. 6E). The core subunit of photosystem psa (D, F, H, L, N, and O) and outer light-harvesting complexes comprising by LHCA (1, 2, and 4) and LHCB (1-5) genes were strongly down-regulated in SWL compared to ST or WL. The ω of LHCB1 (16.33%) LHCA2 (12.24%) LHCA4 (8.16%) were higher compared to the other genes related to light harvest complexes (Fig. 6A). The same pattern was found in genes related to the plastoglobule biosynthesis where the ABC1genes (activity of bc1complex; protein kinase family) and ADCK (aarF domain-containing kinase) were predominant (Fig. 6B). ADCK, ABC1 expression variations are crucial for photosynthetic performance and plant stress tolerance. The polysaccharide catabolic process and carbohydrate transport subset genes showed several genes that were up or down-regulated. The genes ω encoded chitinase (23.81%) beta-amylase (12.7%), endoglucanase (12.7%), and pectinesterase (9.52%) were predominant in carbohydrate catabolism (Fig. 6C). Also, the carbohydrate transporters SLC2A13 related genes showed a considerable high ω in SWL. However, several isoform of SLC2A13 were up or down-regulated (Fig 6D). Fig. 6F-G depicted a dramatic shutdown of the light-harvesting chlorophyll protein complex, photosystem I and II, and the photosynthetic electron transport in *D. odorifera* leaflets under SWL. The down-regulation of photosynthetic-related genes under SWL was intense and quantitatively diverse compared to WL or ST. The glycolysis and citrate cycle were also functional subset genes that could be selected and provide more informative patterns in plant photosynthesis under combined stress. The transcriptional changes in *D. odorifera* leaflets described above resulted in a significant decrease in net photosynthetic rate (Pn) and stomatal conductance (Gs) in SWL compared to control and WL (Fig. 6H-I). However the difference between SWL and ST groups were not significant. The plant hormone variations were more complex, and ABA as well as GA_3_ contents in SWL were statistically enhanced compared to the control (Fig. 6J-K). The IAA concentration was similar among CT, ST, and SWL, but its content in the WL group was lower (Suppl. Fig. 16B).

**Figure 5.**
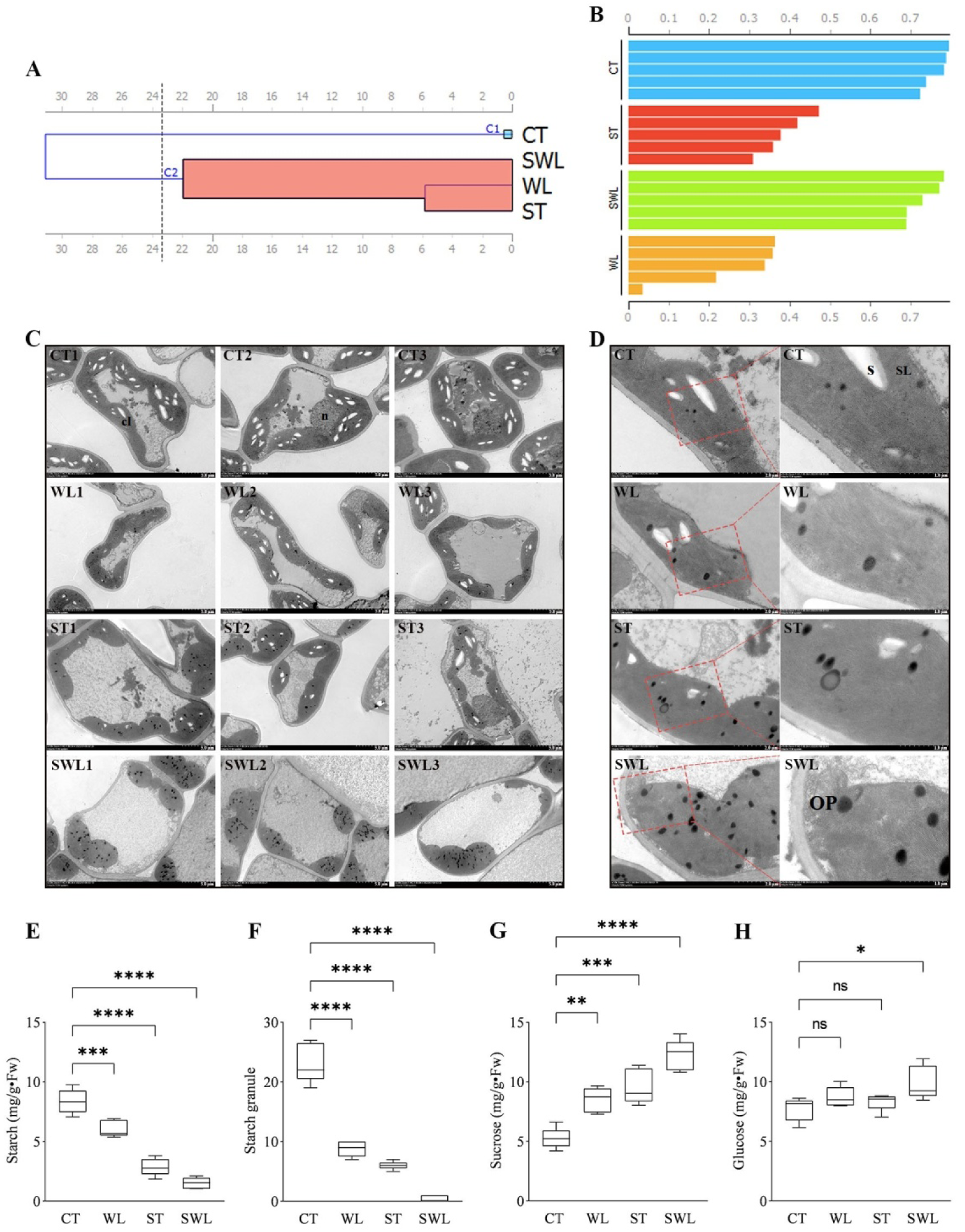
Photosynthesis variations in *Dalbergia odorifera* under single and combined stress, dendrogram of hierarchical clustering of the data set regrouping different photosynthetic related-parameters (glucose, sucrose, and fructose, starch, and number of starch granules, net photosynthetic rate, stomatal conductance, and ABA, GA_3_, and IAA) among CT, ST, WL and SWL (**A**). Silhouette analysis of the different clusters among different treatments based on Euclidean distance metric (**B**). Comparative images of chloroplast ultrastructure that showed starch and plastoglobuli variations among CT, ST, WL and SWL, CL; chloroplast, N; nucleus, OP; osmiophilic plastoglobuli, S; starch granule, SL; stromal lamellae (**C**; 5 µm) (**D**; 2 and 1 µm). Variation of starch content (**E**), the number of starch granules (count visually in TEM pictures) (**E**), sucrose and glucose contents. Treatments are presented as follows: control (CT), salinity (ST), waterlogging (WL), and combination of WL and ST (SWL). The signs such as *, **, ***, **** and NS represented the significance difference among different treatments. *; P < 0.05, **; P < 0.01, ***, P = 0.0001; ****; P < 0.0001, NS; no significant.

**Figure 6.**
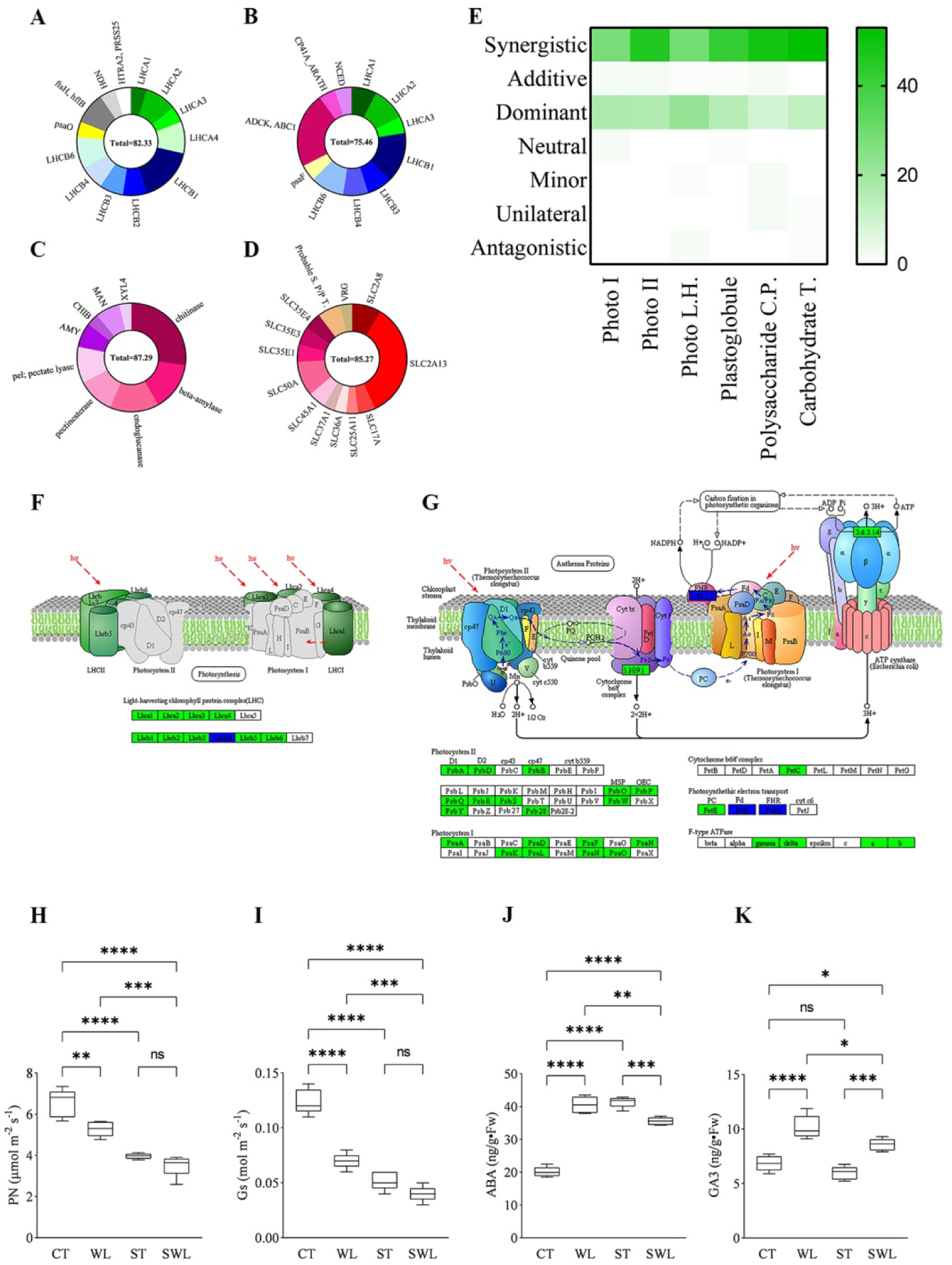
Donut-plots regrouping photosynthesis light harvesting (**A**) plastoglobule (**B**) polysaccharide catabolic process (**C**) and carbohydrate transport (**D**) related genes and their isoforms based on gene ontology (GO) classification and their relative frequency (ω). Heat map of the pattern modes in SWL based on functional GO classification related-DEGs in photosystem I (Photo I), photosystem II (Photo II), photosynthesis light harvesting (Photo L.H.), plastoglobule, polysaccharide catabolic process (Polysaccharide C.P.) and carbohydrate transport (Carbohydrate T.) classes (**D**). Schematic representation of the significantly enriched KEGG pathway diagrams; of the photosynthesis-antenna proteins (**E**) and photosynthetic apparatus(**G**) under SWL that showed the regulation mode and relation of genes belong to the photosystem I and II, cytochrome b6/f complex, photosynthetic electron transport chain and/or F-type ATPase, KO nodes containing only up-regulated differential genes are marked in red, KO nodes containing only down-regulated differential genes are marked in green, and KO nodes containing both up-regulated and down-regulated genes are marked in blue. Variation of net photosynthetic rated (**H**), stomatal conductance (**I**), and ABA (**J**) and GA_3_ (**K**) contents. Treatments are presented as follows: control (CT), salinity (ST), waterlogging (WL), and combination of WL and ST (SWL). The signs such as *, **, ***, **** and NS represented the significance difference among different treatments. *; P < 0.05, **; P < 0.01, ***; P = 0.0001, ****; P < 0.0001, NS; no significant.

### The emphasis on the oxidative transcriptional pattern and physiological modifications under combined stress

Under combined stress, toxic molecules of oxygen accumulation governing plant stress tolerance is a key window to look at and define the pathway in which both single stresses together trigger a global systemic response in oxidoreductase and antioxidant activity. Indeed, over-accumulation of reactive oxygen species (ROS) in plants generated oxidative stress that induced antioxidative mechanisms and redox signaling. The oxidative hierarchical cluster was built based on indexes shown in Suppl. Fig 17-19, the main component comprised ROS, antioxidant enzymes and molecules, proline, phenolic compounds, and related enzymes (PAL and PPO). The items were clustered in two separated groups; ST and SWL and then in WL and SWL (Fig. 7A). Moreover, silhouette analysis showed a cluster in the ST group with a negative value and other clusters with lower values (Fig. 7B) which means that ST antioxidative response was similar to SWL compared to WL. Although for each functional subset of genes, the synergistic effect of combined stress was predominant (Fig. 7C), the biochemical antioxidative response showed a very complex pattern. The transcriptional changes under combined stress showed a high ω of L-ascorbate peroxidase (7.94%), peroxidase (23.81%), and nucleoredoxin (23.81%) related genes and their isoform (Fig. 7D). The gene and its isoform that showed a constant up-regulation status (Suppl. Table 9) were those related to nucleoredoxin which has the function of shield for the antioxidants that are responsible for ROS scavenging. The gene ontology detoxification functional subset genes were mostly represented by GST (glutathione S-transferase) under SWL (Fig. 7F), with a synergistic pattern compared to ST and WL. Most of the GST-related genes were up-regulated, but the concentration of GSH (Suppl. Fig 18C) or the activity of GPX (Suppl. Fig 18F) did not validate the synergistic pattern observed in GST-related transcripts. The oxidoreductase activity functional subset (Fig. 7E) showed a high abundance of transcripts including LOX1_5 (linoleate 9S-lipoxygenase) which had a greater ω compared to the other component such as Laccase, LOX2S, CYP71A, or ABCB1, CD243 (ATP-binding cassette, subfamily B (MDR/TAP), member 1). Furthermore, *D. odorifera* is a medicinal plant with a high phenolic concentration that plays a crucial role in oxidative stress. To highlight the role of flavonoids and other phenolic compounds under SWL, Fig. 7G showed the ω of several genes that were regulated in SWL and their pattern considering their regulation under single stress. The related genes involved in the regulation of phenylpropanoids biosynthesis under SWL that showed a high ω were cinnamyl-alcohol dehydrogenase (9.59%), caffeoyl-CoA O-methyltransferase (10.96%), laccase (9.59%) and 2-hydroxyisoflavanone dehydratase (10.96%). These genes were massively up-regulated (Suppl. Table 9) in SWL compared to ST or WL and followed a synergistic pattern mode. The biochemical analysis showed a high accumulation of the total phenol and flavonoids (Suppl. Fig 19B-C) in SWL, ST, and WL compared to the control. The same pattern was observed in the enzymes such as PAL and PPO (Suppl. Fig 19E-F) involved in phenylpropanoids biosynthesis. Overall, the synergistic pattern spotted in the transcriptional changes in the phenylpropanoids biosynthesis pathway does not reflect the same pattern in the biochemical analysis of the phenolics compounds and enzymes.

**Figure 7.**
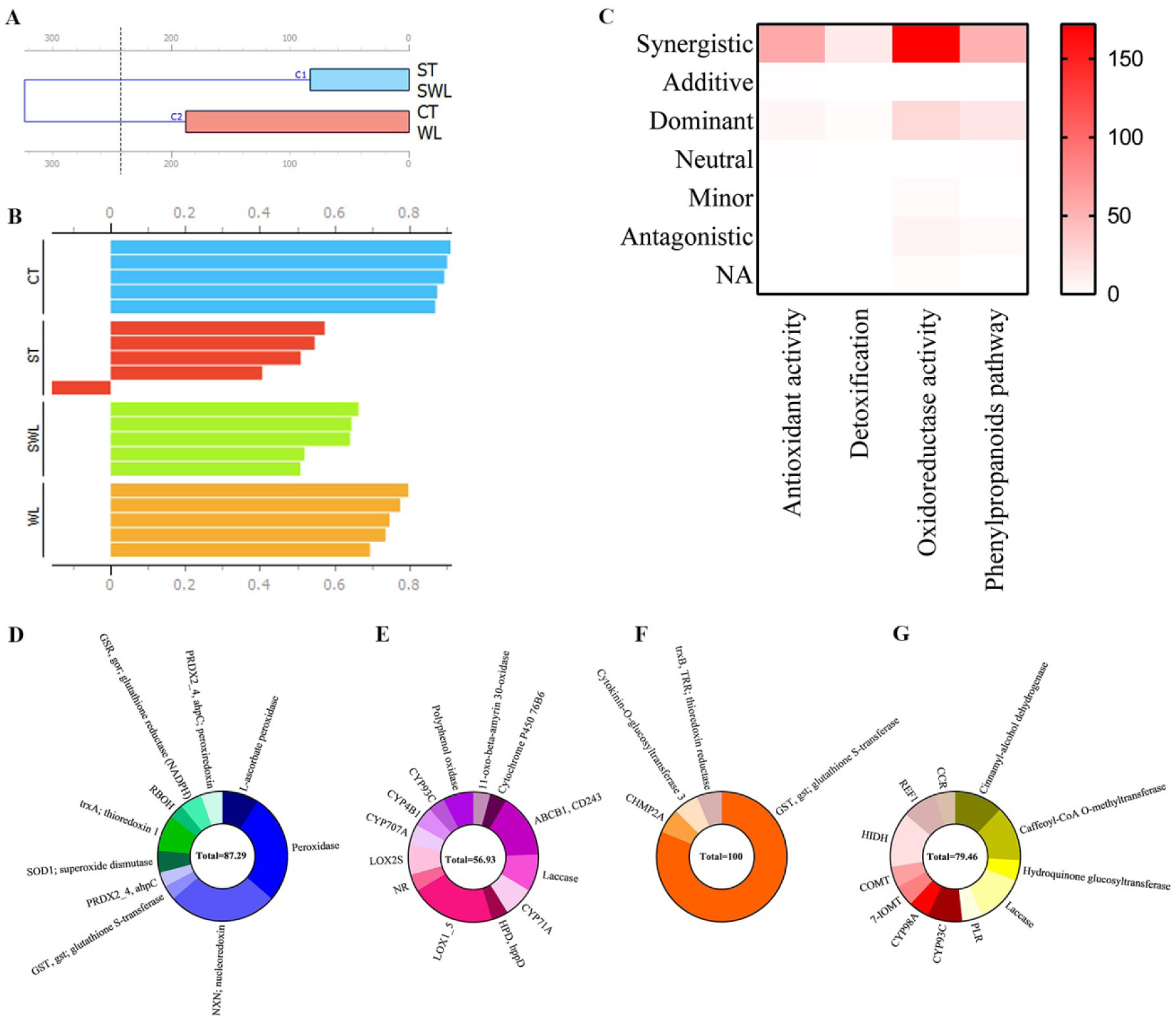
Biochemical and transcriptional profiles of oxidative stress in *Dalbergia odorifera* under single and combined stress, dendrogram of hierarchical clustering of the data set regrouping different oxidative related-indexes (H_2_O_2_, O_2_^•−^, POD, SOD, CAT and GPX, ASA and GSH, proline, phenols, flavonoids, PAL and PPO) among CT, ST, WL and SWL (**A**) and silhouette analysis of the different clusters among different treatments based on Euclidean distance metric (**B**). Heat map of the pattern modes in SWL based on functional GO classification related-DEGs in antioxidant activity, detoxification, oxido-reductase activity and phenylpropanoids pathway classes (**C**). Donut-plots regrouping antioxidant activity (**A**) detoxification (**B**) oxido-reductase activity (**C**) and phenylpropanoids pathway (**D**) related genes and their isoforms based on GO classification and their relative frequency (ω). Treatments are presented as follows: control (CT), salinity (ST), waterlogging (WL), and combination of WL and ST (SWL).

### Does the plant morpho-anatomical analysis validate the metabolomic and transcriptional pattern results?

The anatomical analysis was conducted in healthy leaflet indeed during the first experiment the seedlings that were exposed to a combined WL and 150 mM or 200 mM ST displayed drastic leaflet-damages after one week of treatment as shown in Fig. 8A. Seedlings under WL combined with 100 mM of ST were in a good shape with healthy leaves. The results in this section are in two parts, those taken with *D. odorifera* leaflets and those measured in the roots. We attempt to correlate leaflet and root responses under SWL compared to ST and/or WL and to discover whether the pattern was significantly correlated with the metabolomic and transcriptomic analysis. The hierarchical cluster showed two groups comprised of CT and a second group containing SWL, WL, and ST (Fig. 8B). Two clusters from the ST treatment exhibited a negative silhouette value that brings them closer to SWL compared to those in WL (Fig. 8C). ST and WL presented different types of anatomical changes including large (WL and ST, 1) and small surfaces (WL and ST, 2) of spongy and palisade mesophyll cells (Fig. 8D). The size and organization of the vascular bundle were more consistent in the different samples from CT group. Interestingly, the SWL group showed smaller vascular bundles, spongy and palisade mesophyll cells in all samples observed. The anatomical pictures from each treatment were taken from 6 samples and only 3 samples were presented in this section. At this stage, it’s difficult to determine a pattern in SWL based on ST and WL leaf anatomy observation. To depict the effect of SWL in roots anatomy compared to CT and single stress, the stele that comprised all the tissues inside the endodermis (xylem, phloem, and pericycle) was shown in Fig.8 E. Indeed, Stele is the pivotal part of the root system that contains vascular tissue (xylem and phloem). Seedlings under CT maintained higher (P< 0.05) stele areas (Suppl. Fig 20) than that in SWL treatment which showed the same trend as ST and WL. However, the difference between SWL and ST groups was not statistically significant as well as the difference among ST, WL, and SWL. To further confirm the pattern observed in leaf and root anatomical variation under SWL, we investigated root and leaf morpho-physiology (Fig. 9 and Suppl. Fig 21). The results displayed a strong decrease in shoot height, leaf area, leaf vascular bundle diameter, and shoot water content under SWL (Fig. 9A-D). The root membrane permeability (Fig. 9E) increases significantly (P < 0.05) in ST, WL and SWL. The pattern in SWL was synergistic compared to each single stress. The root water content showed the same trend, and its variation between ST-treated and WL-treated seedlings was similar.

**Figure 8.**
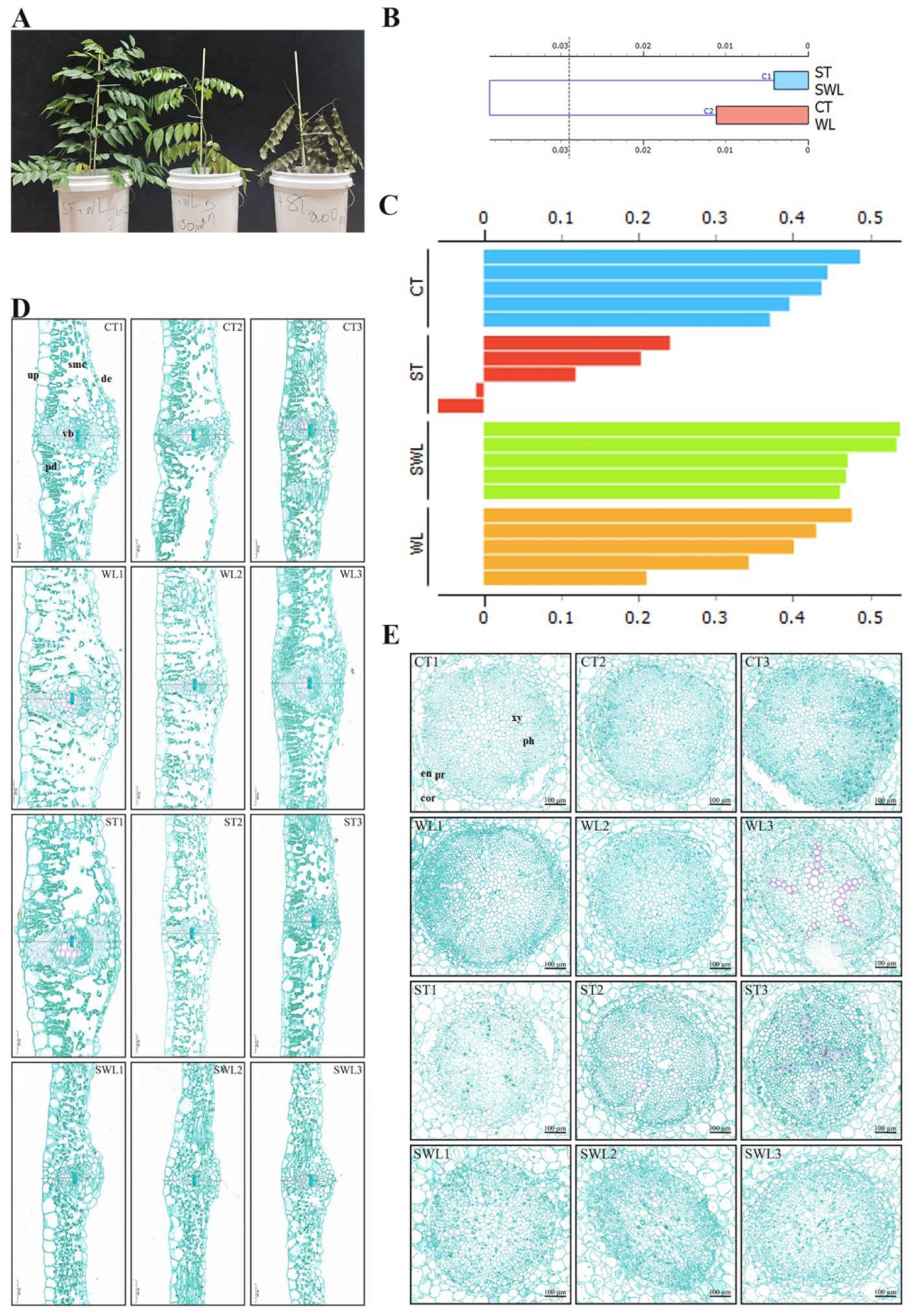
Plant morpho-anatomical responses of *Dalbergia odorifera* under combined stress, photograph of different levels of combined stress (SWL1, SWL2, and SWL3) effects on the shape of the seedlings (**A**). Dendrogram of hierarchical clustering of the data set regrouping different morpho-physiological related-indexes (plant shoot height, shoot and root biomass accumulation, root membrane permeability, shoot and root water contents, leaflet area, leaflet length, leaflet average width, leaflets max width, leaflet vascular bundle diameter and root stele diameter, etc.) among CT, ST, WL and SWL (**B**) and silhouette analysis of the different clusters among different treatments based on Euclidean distance metric (**C**). Anatomical changes of the leaflet (**D**) and root (**E**) of *Dalbergia odorifera* exposed to CT, WL, ST and SWL with 3 replicates each, row 1 (CT1, CT2 and CT3), row 2 (WL1, WL2, and WL3), row 3 (ST1, ST2, and ST3) and row 4 (WL1, WL2 and WL3). Leaflets’ images were taken at a set of 20x and 10x for roots, the zoom level with CaseViewer and scale bars = 50 μm for leaflets and = 100 μm for the roots. The letters within the leaflets images are: de, down-epidermis; pd, palisade mesophyll cell; smc, spongy mesophyll cell; ue, upper-epidermis; vb, vascular bundle.Those in roots images are: cor; cortex, en; endoderme; pr; pericycle, ph; phloem, xy; xylem. Treatments are presented as follows: control (CT), salinity (ST), waterlogging (WL), and combination of WL and ST (SWL).

**Figure 9.**
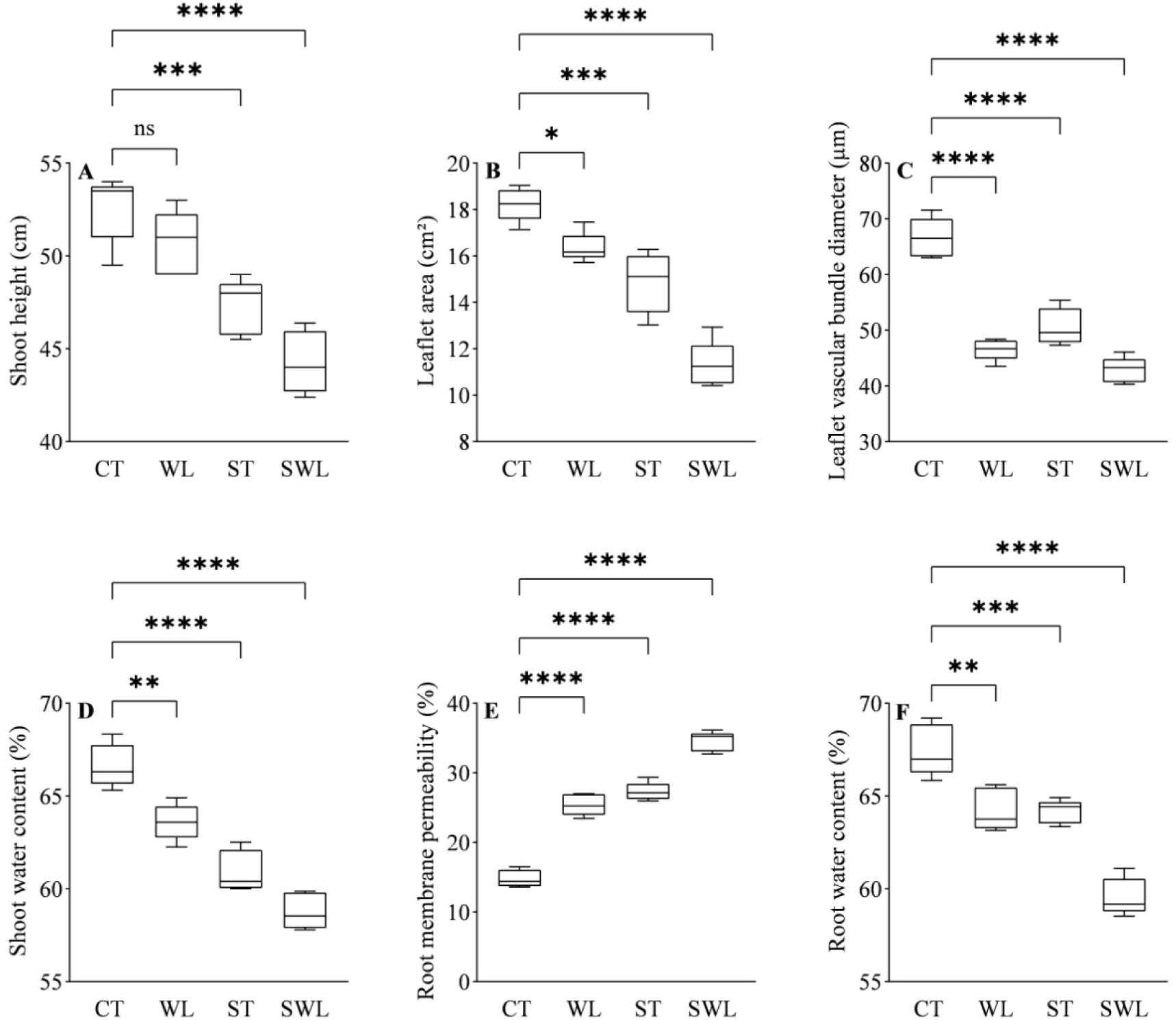
Variation in shoot height (**A)**, leaflet area (**B**), vascular bundle diameter (**C**), shoot water content (**D**), root membrane permeability (**E**) and root water content in *Dalbergia odorifera* seedlings under CT, WL, ST and SWL. Treatments are presented as follows: control (CT), salinity (ST), waterlogging (WL), and combination of WL and ST (SWL). The signs such as *, ***, **** and NS represented the significance difference among different treatments. *; P < 0.05, ***; P = 0.0001, ****; P < 0.0001, NS; no significant.

## Discussion

### Interconnection between multi-factorial stress combination and global climate changes: Challenges and perspectives for woody plant

Global climate change implies an increase in more extreme environmental events that will negatively impact forest tree species distribution and development. Natural catastrophes such as typhoons and storms are making their signature in coastal and wetland areas that influence ecosystem resistance and resilience (Patrick et al., 2022). Trees are one of the main factors that will sustain ecological restoration and management against hazardous floods conditions, which will ipso facto increase the ecosystem resilience (Nilsson et al., 2018). The simultaneous occurrence of waterlogging and salinity is rising in landscapes, and several earlier reports have acknowledged this phenomenon (McFarlane and Williamson, 2002; Carter et al., 2006; Barrett-Lennard and Shabala, 2013). Marcar et al., (2002) established that commercial of seeds, seedlings or tree species depended on multi-factorial stress tolerance, particularly with the increasing event-related to climate change. Here comes advocacy about the scarcity of molecular studies in tree species compared to crops. Sustain food security through comprehensive research focused on molecular profile and pathway in crops under harsh conditions is one imperative thing to achieve. But, can it be imagined to reach that objective without an equal understanding of the molecular response of the species that sustain most of the terrestrial ecosystems? Emergence of significant research on plant molecular responses under combined stress conducted mainly in crops or *Arabidopsis* has prompted the scientific community to realize that plant response to single stress is remarkably distinct compared to multi-factorial stress conditions. Thus it is worthwhile to explore the molecular responses integrated with physiology of woody plants under simultaneous waterlogging and salinity. Surprisingly the transcriptional changes integrated with physiological and metabolomic profiles in plants under hypoxia combined with salinity are quasi-inexistent.

### Combined stress increased quantitatively and qualitatively the number and level of transcripts and metabolites following various pattern modes

The molecular profiles of plant species subjected to various combinations of environmental stress are uncommon to the responses from each individual stress (Zandalinas and Mittler, 2022). The combinatory stress among salinity, drought, and heat, cold or high light revealed a smaller percentage of similarity found in transcript expression compared to single stress. The metabolomic changes showed the same pattern with an accumulation of various non-structural carbohydrates presents only in plants under combined stress (Mittler, 2006). The results of the present study showed an impressive percentage of increase in transcripts expressed under SWL compared to WL (241.8%) or ST (426.3%). The increase in number of transcripts correlated positively with gene enrichment following their biological functions. The combination of high light and heat stress in *Arabidopsis* showed a similar trend; indeed, GO term enrichment analysis provided remarkable transcriptional responses that included genes related to photosynthesis, oxidoreductase activity, or carbon and protein catabolism and transport (Prasch and Sonnewald, 2013; Shaar-Moshe et al., 2017; Balfagón et al., 2019). Guo et al., (2021) found a massive enhancement of amino acids in maize during combined drought and cold stress. The uniqueness of metabolomic response in *D. odorifera* leaflets under SWL (Suppl. Table 4-5) was based on the repression of lipids such as glycolipids comprising monogalactosyldiacylglycerol (MGMG [18:20] isomer2) and digalactosyl-diacylglycerol (DGMG [18:1]). The up-regulation of these glycolipids under WL or ST would be related to their stress-related function in membrane stabilization and stress signaling (Zhang et al., 2019). Among the lipids, monoacylglycerols (MAG [18:3] isomer5), 9-Hydroxy-(10E, 12Z, 15Z)-octade-catrienoic acid and 13-hydroxy-9, 11, 15-octadecatrienoic (13-HOTrE) were also uniquely down-regulated under SWL. Moreover, the biosynthesis of several amino acids has been triggered by SWL and not through ST or WL. Phenylalanine, tyrosine, aspartic acid, pyrrole-2-carboxylic acid, histidine, aminoadipic acid, aspartic acid, and methionine were up-regulated by SWL while they were suppressed in WL or unchanged under ST. SWL showed an antagonistic effect in amino acid accumulation in *D. odorifera* leaflets. The accumulation of amino acids in plant under harsh conditions is a requirement for an alternative respiratory substrate (Batista-Silva et al., 2019). The introduction above of the metabolomic variations in *D. odorifera* leaflets referring to the lipids repression and amino acid accumulation as a specific response of combined stress compared to ST or WL, supports and validates the need for determination of metabolites and genes pattern of plants under combined stress conditions. The method described in the methodology section (Expression Patterns of genes and metabolites) has been influenced by earlier studies focused on the transcriptional response of *B. dystachion* under combined salinity, drought, and heat (Shaar-Moshe et al., 2017). Nevertheless, new and different approaches of genes and/or metabolites pattern provided by the present study confirm an unavoidable method to depict, demonstrate and discover the uniqueness of molecular response of combined stress in plant following their biological function and/or classification. Indeed, the results showed 7.38% of the metabolites in the antagonistic mode, among them, 60.9% were classified as lipids, and the second most abundant class was represented by the amino acids (12.1%), which demonstrated the uniqueness in common metabolites regulation under combined stress. The suppl. Table 8 and 9 showed several unique genes and their isoform following GO classification that were regulated under combined stress which was attributed to a synergistic mode. The down-regulation of DOPA (4,5-DOPA dioxygenase extradiol) related genes under SWL or the up-regulation of FMO (dimethylaniline monooxygenase [N-oxide forming]) were both in synergistic mode compared to a single stress. Their regulation might be depending only on a combinatory stress mode between ST and WL. Moreover, a closer look at the different common transcripts pattern modes among ST, WL, and SWL showed that several genes related to response to stress, acyl-CoA oxidase (*ACOX1*, *ACOX3*), *FAD2* (omega-6 fatty acid desaturase) or *ASNS* (asparagine synthase) presented a synergistic effect in SWL. The suppl. Table 11 summarized how combined stress synergistically affects the genes mainly expressed in plants under stress. GO classification showed several genes from SWL-treated seedlings that are in the module Response to Stress, including the molecular chaperone HtpG, trehalose 6-phosphate synthase/phosphatase (*TPS*), mitogen-activated protein kinase 3 (*MPK3*), and calcium-permeable stress-gated cation channel (*TMEM63*) showed a massive expression compared to ST or WL.

Based on common transcript pattern modes classification, it is possible to quickly isolate and identify the genes or metabolites crucial in plant tolerance under combined stress. Providing a transcriptional and metabolomic pattern mode in plants under combined stress is a significant methodology that can be materialized by bioinformatics program packages that generate diagrams, scatter plots, or heat maps. Indeed, the genes and metabolites expression pattern in the present study has been done manually. The development of bioinformatics tools such as an R package program could change how we generate data from combined stress and help the scientific community isolate several genes key to improving stress tolerance in plants under combined stress.

### Shut down of the photosynthetic apparatus through a massive down-regulation of *LHCBs* and *PSAs* genes. A less-needed role of ABA in signaling pathway under SWL?

The data showed a massive synergistic shutdown in the photosynthetic apparatus reflected by the suppression of several light-harvesting complexes (I and II) chlorophyll a/b binding proteins, photosystem I subunits (II, III, VI, XI, *PsaN*, and *PsaO*) related genes in PSI. The PSII as a multi-subunit enzymatic complex that catalyzes the breakdown of water molecules during photosynthesis (Rhee et al., 1998) showed the same pattern as PSI. The high frequency of repression of *LHCB1*, *LHCA2*, *LHCA4*, *LHCB1*, *LHCB2*, *LHCB3*, *LHCB4*, and *LHCB6* in PSII was correlated with a significant decrease in ABA content. Over-expression of the *LHCB6* gene in *Arabidopsis* induced ABA accumulation that favoured stomatal response (Xu et al., 2012). It was suggested that the ABA receptor that triggers downstream signaling cascades to induce photosynthetic changes is strongly dependent on the regulation of *LHCBs*. Indeed, over-expressed *LHCBs* in transgenic *Arabidopsis* showed an up-regulation of two transcription factors (*ABF2* and *ABF3*) that are positively involved in ABA signaling (Xu et al., 2012). *ABF* transcripts were more up-regulated in *D. odorifera* leaflets under SWL compared to ST or WL (Suppl. Table 10) and along with *ABA1* (zeaxanthin epoxidase) and *ABA3* (molybdenum cofactor sulfurtransferase) genes; 9-cis-epoxy carotenoid dioxygenase (*NCED*) gene was repressed under combined stress and induced under ST or WL. However, *ABA2* genes showed an increased level of expression in SWL-treated seedlings similar to ST-treated seedlings. A previous report mentioned that *ABA1*, *ABA3*, and *AAO3* (abscisic-aldehyde oxidase) related genes might be more sensitive to the ABA-mediated positive feedback on ABA biosynthesis (Xiong et al., 2002). This would explain a part of the mechanistic molecular pathway behind the decrease of ABA content compared to ST or WL (Fig. 6J) in *D. odorifera* leaflets triggered by SWL. Indeed, as shown in Suppl. Table 10 *AAO3* was significantly induced by SWL, but its activity required a sulfurated molybdenum cofactor (MoCo) and the conversion of MoCo pass by *ABA3* (Barrero et al., 2006 and references therein) which was down-regulated in SWL-treated seedlings. The results highlighted that ABA1 might have a more important function compared to *ABA2* or *ABA3* in ABA regulation under SWL. It has been well-established that *ABA1* genes catalyze the epoxidation of zeaxanthin to antheraxanthin and all-trans-violaxanthin which is converted into the cis-isomers of neoxanthin and violaxanthin cleaved by *NCED* to form xanthoxin. Xanthoxin is converts into abscisic aldehyde which is an intermediate product in the biosynthesis of the abscisic acid (Perreau et al., 2020; Qin & Zeevaart, 1999; Schwartz et al., 1997). The positive correlation between *ABA1* and *NCED* would suggest similar importance of both genes in the decrease of ABA level compared to single stress in *D. odorifera* leaflets under SWL. Although the ABA level was significantly lower in SWL-treated seedlings than in ST or WL, surprisingly the net photosynthetic rate and stomatal conductance variations were lower in SWL compared to ST or WL but significantly greater than that in CT. Previous research found a synergistic accumulation of ABA in maize under cold associated with drought (Guo et al., 2021) or in *Arabidopsis* subjected to simultaneous Heat and High light (HHL) (Balfagón et al., 2019). *Arabidopsis* ABA mutant deficient did not show a high sensitivity to HHL compared to the wild-type, thus ABA showed a lesser role compared to other plant hormones such as jasmonic acid in HHL tolerance (Balfagón et al., 2019). However, Guo et al., (2021) demonstrated that ABA is crucial to maize adaptation to combined cold and drought (CD). Moreover, the stomatal conductance and net photosynthetic rates in maize were disproportional to the massive increase of ABA in CD compared to the single stresses. Indeed, between drought and CD-treated seedlings the stomatal conductance was statistically similar (Guo et al., 2021) which would suggested other roles of ABA in CD-tolerance. Our results suggested that under combined stress a synergistic accumulation of ABA in plant leaflets is not required to regulate the stomata and net photosynthetic rates. In addition, the up-regulation of several transcripts that encoded PSI (PsaA, PsaK, PsaC, or PsaH) and PSII (PsbC, PsbA, PsbB, PsbE, PsbF, PsbH, or PsbZ) proteins in *Arabidopsis* under HHL (Balfagón et al., 2019) was contradictory to the results found in *D. odorifera* (Suppl. Table 8). SWL triggered a strong repression of an impressive number of photosystem subunit related genes and isoform in PSI (*psaA*, *psaF*, *psaD*, *psaL*, *psaH*, *psaN*, *psaO*, and *psaK*) and PSII (*psbQ*, *Psb*, *psbO*, *psbB*, *psbR*, *psbW*, *psbY*, *psbA*, *psbS*, *psbD*, and *psbP*).

### Starch degradation and plastoglobuli synthesis emerged to play a crucial role in *D. odorifera* chloroplast under SWL

The shutdown mode of the photosynthesis apparatus by SWL was also applied to *ABC1* genes (*ADCK*; aarF domain-containing kinase), and *ABC1* genes’ loss-of-function in transgenic plants strongly weaken the photosynthetic performance and stress tolerance in plants (Espinoza-Corral and Lundquist, 2022). Several ABC1 proteins were identified in *Arabidopsis* chloroplasts, specifically in the plastoglobuli proteome (Lundquist, Poliakov, et al., 2012), Martinis et al., (2013) suggested a stabilizing function of ABC1-like kinase ABC1K3 in Arabidopsis plastoglobuli. The repression of *ABC1* genes under SWL correlated with a remarkable increase of osmiophilic plastoglobuli in *D. odorifera* chloroplast that would confirm the stabilizer function of the ABC1 proteins in plastoglobuli formation. It is well-known that abiotic stress affects plastoglobuli (lipoprotein bound to thylakoid) morphology which is closely related to their composition (Arzac et al., 2022). The proliferation of osmiophilic plastoglobuli in *D. odorifera* chloroplasts under SWL was chaperoned by a spectacular degradation of starch granules (Fig. 5C-D). Glucose, sucrose, and fructose accumulation were higher in SWL-treated seedlings resulting in starch degradation. The metabolomic analysis showed a considerable up-regulation of sucralose in *D. odorifera* leaflets under SWL, ST, or WL. The main factors that trigger the repression of starch accumulation in SWL were related to the down-regulation of *WAXY* (granule-bound starch synthase) and *glgA* (starch synthase) genes (Suppl. Table 10). Stress-induced starch degradation is a well-known pathway in plant stress tolerance acquisition and improvement (Thalmann and Santelia, 2017). Moreover, a specificity of SWL was the suppression of genes encoding pectinesterase and endoglucanase, but several genes and their isoform related to chitinase and beta-amylase were strongly induced by SWL (Suppl. Table 8). The *β*-amylase is one of the main enzymes in starch degradation; β-amylase deficient-of-function in transgenic potato leaves indicated a defect in starch degradation (Scheidig et al., 2002). Starch is a polymer of sugar formed in chloroplasts and represents the most widespread glucose-based reserve in plant leaves; it is synthesized through starch synthase during photo-assimilated CO_2_ during photosynthesis (Bürgy et al., 2021; Szydlowski et al., 2009). However, plants under abiotic stress-induced starch degradation reduce their photosynthetic activity. Amylose as a major component of starch is synthesized through granule-bound starch synthase (Lorberth et al., 1998). Hypoxia combined with different ST concentrations showed to improve sugar monomers accumulation in *Suaeda maritime* compared to single stress conditions (Behr et al., 2017). The same pattern was found in maize under combined cold and drought in maize (Guo et al., 2021) or drought and heat in *Arabidopsis* (Prasch & Sonnewald, 2013). Further, the role of sugar transporter under combined stress is largely unexplored. However, a significant increase of the monomer facilitated transporter such as SLC2A13 (MFS transporter, SP family, solute carrier family 2 (myoinositol transporter), member 13) was noticed under SWL compared to ST, WL, or CT. The results suggested a crucial function of sugars transport in *D. odorifera* leaflet under SWL.

### Nucleoredoxin-related transcripts (*NXN*) maintained the high synergistic antioxidative system in *D. odorifera* leaflets under SWL and cross-talks among oxidative stress, antioxidant and phenylpropanoids

Oxidative stress is rarely discussed in the small number of scientific papers that have studied the transcriptional responses of combined stress on plants. Oxidative stress is a primary reaction of plants cells under stress triggered mainly by an accumulation of toxic molecules of oxygen known as reactive oxygen species (ROS). Knowing the changes in oxidative status in plants under combined stress is crucial due the fact that ROS are mainly produced in the chloroplasts and peroxisomes through photorespiration (Waszczak et al., 2018). The lifetimes of hydrogen peroxide (H_2_O_2_) and superoxide anions (O_2_^•−^) that are considerably longer (milliseconds to seconds) compared to other ROS (Sies et al., 2017) were measured and showed a strong accumulation in SWL-treated leaflets. The over-production of H_2_O_2_ and O_2_^•−^ in *D. odorifera* under ST, WL, or SWL is related to various functions of ROS in stress tolerance such as signaling pathways or programming cell death. ROS-induced DNA, lipids, or proteins damages are related to a defective functioning of its detoxification (scavenging or quenching) (Foyer and Hanke, 2022). The strong up-regulation of ROS-response transcripts in Arabidopsis described by Zandalinas et al., (2020) under a combination of high light and heat stress showed a significant key role of ROS in plant systemic signals and responses under combined stress. But still, these results highlighted only the function of ROS in plant signaling response under stress integrated with salicylic acid-response transcripts. An earlier study showed two osmolytes (proline and glycine betaine) that have a key role in ROS-scavenging in tomato plants under salinity combined with heat stress. But ROS scavenging under abiotic stress is more complex; two major antioxidative systems are implicated in ROS scavenging; an enzymatic (POD, SOD, CAT or GPX) and nonenzymatic component (ASA or GSH) (Sies et al., 2017). Here, it has been found a dual regulation of peroxidase and L-ascorbate peroxidase-related transcripts (up and down-regulation) under combined stress, and most of the glutathione S-transferase-related transcripts were up-regulated. One of the main factors that lead the antioxidant system defense against oxidative stress in *D. odorifera* leaflets under SWL was the impressive number of nucleoredoxin-related transcripts up-regulated (NXN). In Arabidopsis plants, Kneeshaw et al., (2017) exposed a determinant function of *NRX1*, a member of the TRX superfamily of enzymes in maintaining catalase enzymes in a reduced state thereby protecting their H_2_O_2_-detoxifying activity which ipso facto safeguarded the antioxidant system during oxidative stress. The content of AOX proteins, GSH and ASA increased in SWL along with POD, SOD, and GPX which would confirm an efficient protective role of NXN-related genes of the antioxidant system. *AOX1* and *AOX2* (ubiquinol oxidase) transcripts along with AOX proteins were up-regulated in SWL-treated seedlings. *AOX1* has shown critical function in maintaining the chloroplastic redox state in *Arabidopsis* under abiotic stress by modulating cellular redox homeostasis and ROS accumulation when electron transport through the COX pathway is disturbed at complex III (Vishwakarma et al., 2015). Further, cross-talks between oxidative stress and phenylpropanoid pathways in combined stress are scarce. *D. odorifera* is a medicinal with phenolic compounds present in its organs; it has been noticed that environmental stress can improve the phenolic compounds profiles and activities in plant (Dixon and Paiva, 1995).The content of phenols, flavonoids, and enzymes involve in the phenylpropanoids pathway (PAL and PPO) were increased in ST or LW stress and SWL, but there was not a synergistic pattern in the biochemical and metabolomic characterization of phenylpropanoid pathways. However, there were synergistic transcriptional responses of *D. odorifera* leaflets under SWL compared to a single stress. The non-reflection of the transcriptional response into the biochemical and metabolic characterization under SWL can be explained mainly by the down-regulation of several gene keys and their isoforms involved in the phenylpropanoid pathways such as laccase-related genes. The caffeoyl-CoA O-methyltransferase, cinnamyl-alcohol dehydrogenase, and 2-hydroxyisoflavanone dehydratase-related transcripts were mainly up-regulated in SWL with a synergistic mode compared to a single stress. Laccase-related genes were up- and down-regulated under SWL and were unchanged in single stress. Although caffeoyl-CoA O-methyltransferase, cinnamyl-alcohol dehydrogenase, and 2-hydroxyisoflavanone dehydratase-related transcripts were strongly up-regulated, the down-regulation of laccase enzyme would suggest the non-synergistic effect of SWL in *D. odorifera* leaflets in phenols accumulation.

### The non-synergistic pattern in ABA accumulation would suggest an increase in different other signaling pathway in *D. odorifera* under SWL

ROS and ABA are well-known to play crucial in plant signaling response against osmotic stress mainly triggered in plants by abiotic stresses through the regulation of a broad spectrum of osmotic stress-responsive genes (Sies et al., 2017; Zhao et al., 2022). The ROS up-regulation under SWL was more important compared to ABA which showed a lower concentration compared to ST or WL. That would suggest a less role of ABA in signaling response in *D. odorifera* seedlings under SWL. Plant MAPK signaling pathway map revealed (Suppl. Fig. 22) more genes induced by SWL involved in H_2_O_2_ or ethylene signaling pathways compared to ABA in stress tolerance responses or adaptation. Several plant MAPKs are triggered by a variety of abiotic stress (Zhang and Klessig, 2001), the results would suggest a more important function of MAPKs and ROS in SWL-signaling response. Although ABA content and ABA-related genes showed a correlation in the reduction of ABA activity in *D. odorifera* leaflets under SWL, plant hormone signal transduction (PSL)-related genes showed significant enrichment (Suppl. Fig. 5). The IAA and GA_3_ content did not massively increase under SWL, thus it seems others plant hormones such as ethylene as suggested the MAPK map would play a crucial role in SWL-tolerance. Toll-like receptors, NF-kappa B, neurotrophin, and along with PSL signaling pathways showed a massive role in SWL-signaling responses.

## Conclusion

Plant facing combined stress seems to reveal a general theory which is based on a synergistic transcriptional response due to two factors (i) the stress intensity received by the plants which is related to the simple addition of one stress to another one (ii) and the integration of each specific response related to the specific stress tolerance mechanism of each single stress taking part in the stress combination. In this study, we deciphered experimentally through shock stress different molecular mechanisms involved in SWL in *D. odorifera* seedlings. Combined stress triggered an impressive number of transcripts and differential metabolites compared to single stress. The main biological functions discussed in the present study displayed synergistic responses of *D. odorifera* leaflets under SWL at some points. The analysis of individual transcript or metabolite patterns showed seven different modes where the neutral mode was the most represented. The focus on the biological function of metabolites and transcripts showed that the synergistic effect of SWL is not a general rule in plants following the biological function or type of parameters studied. ABA which is a crucial plant hormone related to stress tolerance and signaling pathway was decreased in SWL-treated seedlings compared to ST or WL. The down-regulation of *ABA*-related transcript *ABA1* or *NCDE* showed the same trend. Nevertheless, the ABA was significantly higher in SWL compared to the control group which partly permits a low net photosynthetic and stomatal conductance variation in SWL-treated seedlings. Indeed the repression of *LHCBs* and photosystem subunit-related genes and isoforms (*psbs*) in PSI and PSII triggered by SWL appear to be the main factor in the shutdown of the photosynthetic apparatus. The massive degradation of starch into monomers of sugar through *β*-amylase along with the increase in sugars transporter such as *SLC2A13* characterized dynamic coordination in the response of *D. odorifera* leaflets to SWL. The same trend was also shown in the elaboration of antioxidative system defence against oxidative stress. Collectively, the biomass accumulation, water content and height of the shoot, leaf area, root anatomy, membrane permeability, and water content validated the transcriptional variation in *D. odorifera* under SWL. Knowledge is changing so quickly and climate change increase so quickly that the scientific community is urged to provide a meaningful molecular database to increase every new concept in plant stress tolerance, thus making each contribution a bit like a grand experiment.

## Acknowledgments and Funding

This work was supported by the National Natural Science Foundation of China (no. 32060240 and 31660165), the Hainan Provincial Natural Science Foundation of China (421RC1033), and Hainan Province Science and Technology Special Fund (ZDYF2022SHFZ054).

## Author contributions

Yang F and Cisse EHM wrote the draft manuscript and draw the graphs, analyzed and interpreted the data; Cisse EHM and Jiang BH performed the most of the experiments and collected the data; Yin L-Y and Miao L-F managed the whole experiments and performed the partial experiments; Li DD, Xiang SL, Mekontso FN, and Zhou JJ assisted in carrying out the partial experiments and data collection; and Yang F designed the experiments, revised the manuscript, and provided funding.

## Ethics approval and consent to participate

*Dalbergia odorifera* T.C. Chen belongs to protected plant species; the trade of its commercialized seedlings is permitted and legal in China. Thus, we settled that no specific permissions for the seedlings collection in this location were required by the Forestry Bureau of Hainan Province, China.

